# A critical but divergent role of PRDM14 in human primordial germ cell fate revealed by inducible degrons

**DOI:** 10.1101/563072

**Authors:** Anastasiya Sybirna, Walfred W.C. Tang, Sabine Dietmann, Wolfram H. Gruhn, M. Azim Surani

## Abstract

PRDM14 is a crucial regulator of mouse primordial germ cells (mPGC), epigenetic reprogramming and pluripotency, but its role in the evolutionarily divergent regulatory network of human PGCs (hPGCs) remains unclear. Besides, a previous knockdown study indicated that PRDM14 might be dispensable for human germ cell fate. Here, we decided to use inducible degrons for a more rapid and comprehensive PRDM14 depletion. We show that PRDM14 loss results in significantly reduced specification efficiency and an aberrant transcriptome of human PGC-like cells (hPGCLCs) obtained *in vitro* from human embryonic stem cells (hESCs). Chromatin immunoprecipitation and transcriptomic analyses suggest that PRDM14 cooperates with TFAP2C and BLIMP1 to upregulate germ cell and pluripotency genes, while repressing WNT signalling and somatic markers. Notably, PRDM14 targets are not conserved between mouse and human, emphasising the divergent molecular mechanisms of PGC specification. The effectiveness of degrons for acute protein depletion is widely applicable in various developmental contexts.

## Main

Gametes develop from primordial germ cells (PGCs), the embryonic precursors, which are apparently specified in ~week 2-3 (Wk2-3) human embryos^1, 2^. While the requirement for BMP and WNT signalling for PGC specification is conserved between mouse and human^3–5^, the gene regulatory network for hPGC fate has diverged^6^. Mouse PGCs (mPGCs) are specified by three core transcription factors (TFs): *Prdm1* (encoding BLIMP1), *Prdm14* and *Tfap2c* (encoding AP2γ)^7, 8^, among which PRDM14 plays a central role; loss of *Prdm14* abrogates mPGC specification^9^, while its overexpression is sufficient to induce mPGC fate *in vitro*^8^.

During mPGC specification, PRDM14 induces upregulation of germline-specific genes, assists BLIMP1-mediated repression of somatic transcripts and initiates global epigenetic reprogramming^7, 8, 10, 11^. PRDM14 also has a significant role in preimplantaion development^12^, as well as pluripotency induction and maintenance in both mouse and human^13–16^. Indeed, *PRDM14* knockdown in hESCs led to a decrease in OCT4 levels and elevated expression of lineage markers^13, 17, 18^.

Despite its critical function in mPGC specification, the role of PRDM14 in hPGC development remains uncertain, due to its low and potentially cytoplasmic expression in gonadal hPGCs^3^. Furthermore, a partial *PRDM14* knockdown suggested it might not be important for hPGC specification *in vitro*^19^, within the TF network for hPGC specification that has diverged significantly from mouse^1, 6, 20^. In particular, SOX17 is a key determinant of hPGC fate, acting upstream of BLIMP1 and TFAP2C^3^, but it is dispensable for mPGC development^21, 22^. Understanding whether PRDM14 has a role in hPGC specification is critical towards gaining insights on the molecular divergence between mouse and human PGCs.

An inducible system for PRDM14 loss of function during hPGCLC specification from hESCs is critical, since PRDM14 is also vital for hESC pluripotency^13^. Accordingly, we combined auxin-or jasmonate-inducible degrons^23, 24^ with CRISPR/Cas9 genome editing^25^ to achieve fast, comprehensive and reversible loss of endogenous PRDM14 protein. We reveal an indispensable role for PRDM14 in germ cell fate, since loss of function affects the efficiency of specification and results in an aberrant hPGCLC transcriptome. Notably, PRDM14 targets are not conserved between mouse and human, reflecting the evolutionary divergence in the molecular network for PGC specification. The study also illustrates the power of conditional degrons, which can be widely used to study TFs during cell fate determination.

## Results

### Detection of dynamic PRDM14 expression during hPGCLC specification

To follow PRDM14 expression during hPGCLC specification, we appended Venus fluorescent protein to the C-terminus of endogenous PRDM14 (Fig.1A) in the background of NANOS3-tdTomato hPGCLC-specific reporter^5^. PRDM14-T2A-Venus line served for flow cytometry and fluorescence-activated cell sorting (FACS) of PRDM14+ cells (Fig.1B-C), while the fusion PRDM14-AID-Venus reporter was used to confirm subcellular localisation of PRDM14 (Fig.1E), as well as for inducible protein degradation (see below). We detected Venus fluorescence in targeted hESCs and hPGCLCs but not in the parental control (Fig.1B-C). Immunofluorescence (IF) confirmed co-localisation of Venus and PRDM14 in nuclei of both hESCs and hPGCLCs (Fig.1E, Fig.2A). Importantly, the majority of alkaline phosphatase (AP)^+^NANOS3-tdTomato^+^ hPGCLCs were PRDM14-Venus+ (Fig.1C) and Venus+AP+ cells specifically expressed key germ cell markers (Fig.1D).

**Figure 1.**
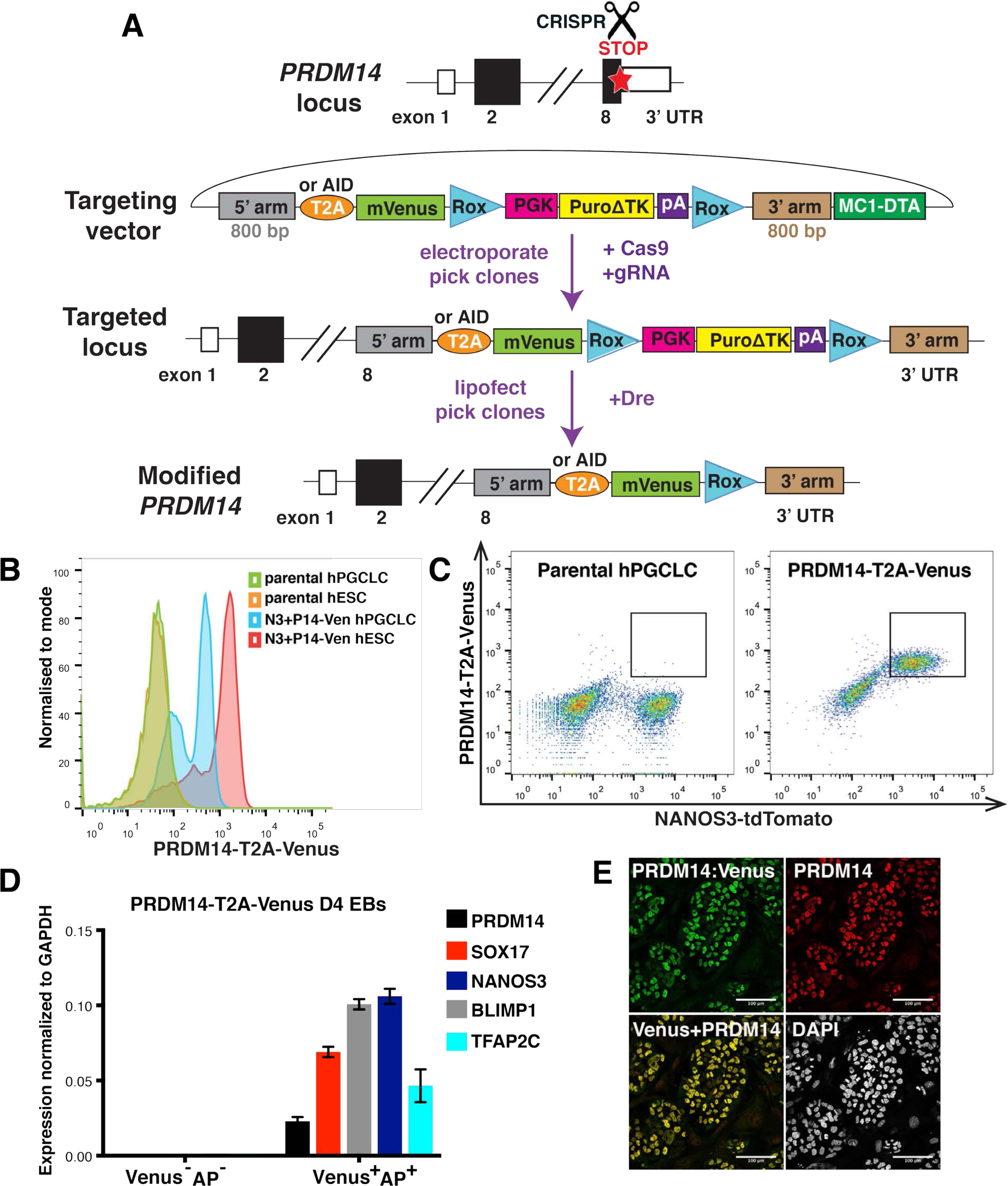
PRDM14-Venus knock-in reporters allow PRDM14 detection in hESCs and hPGCLCs. (**A**) Scheme of CRISPR/Cas9-mediated *PRDM14* locus targeting to generate T2A-Venus or AID-Venus (fusion) reporter versions. 5’ and 3’ arms – homology sequences, T2A – self-cleaving peptide, AID – auxin-inducible degron, Venus – fluorescent gene, Rox – sequences for site-specific recombination recognized by the Dre enzyme, PGK-Puro – puromycin resistance gene under the control of PGK promoter, ΔTK – truncated thymidine kinase gene, MC1-DTA – diphtheria toxin fragment A gene under the control of MC1 promoter. (**B-C**) Flow cytometry analysis showing Venus fluorescence in targeted hESCs and hPGCLCs compared to negative control. Note that Venus fluorescence predominantly coincides with NANOS3-tdTomato signal, which marks hPGCLCs. (**D**) qPCR analysis on sorted PRŋM14-T2A-Venus+AP+ and double-negative cells from D4 EBs. Venus^+^AP^+^ population shows specific expression of germ cell markers. Data show mean+/-SD of 3 technical replicates, n=1. (**E**) IF analysis of PRDM14-AID-Venus in competent hESCs showing colocalization of PRDM14 and Venus fluorescence. Nuclei were counterstained by DAPI. Scale bar is 100 μM.

**Figure 2.**
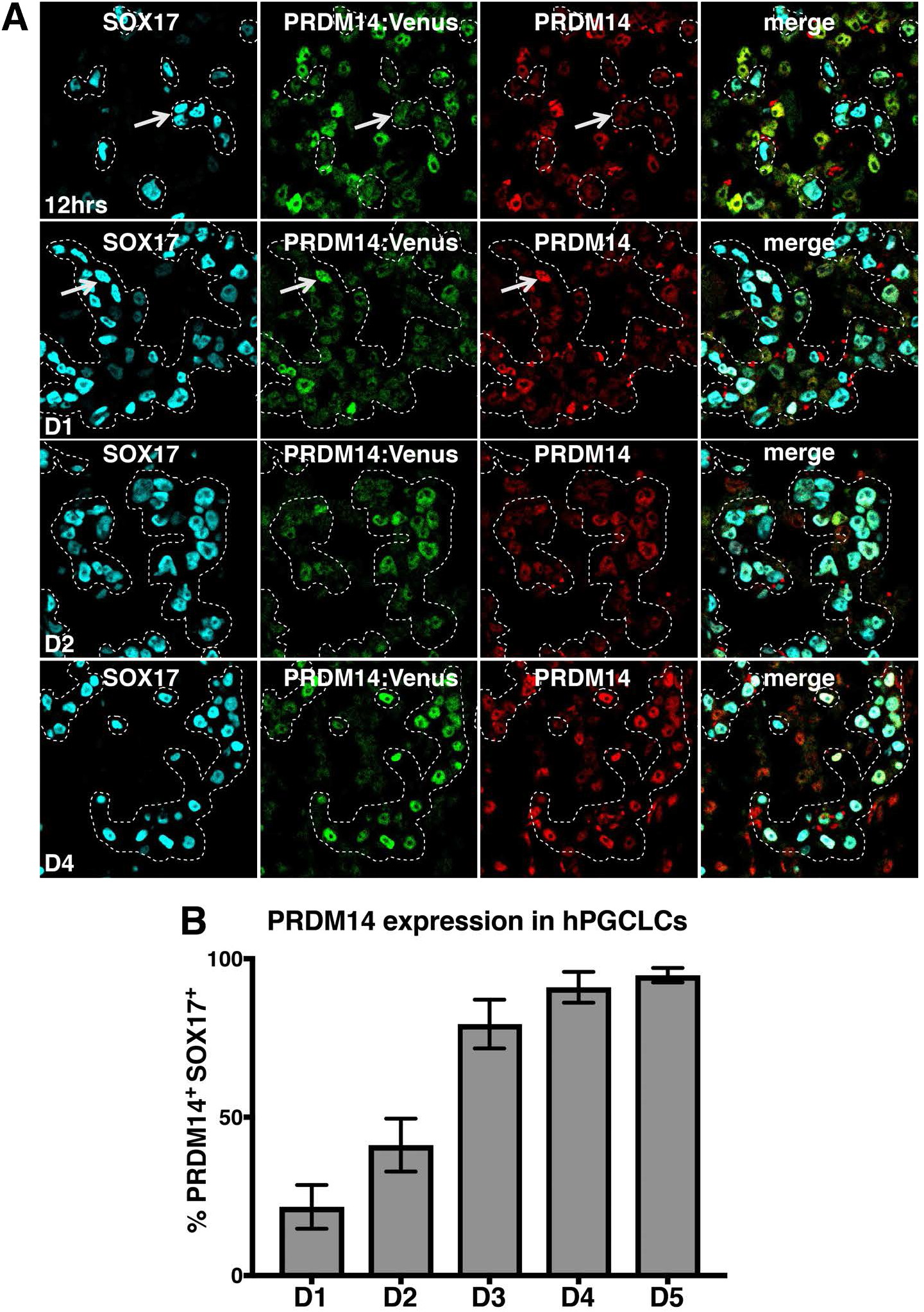
PRDM14-Venus fusion knock-in reporter detects the dynamics of specific PRDM14 expression in hESCs and hPGCLCs. (**A**) Time-course IF analysis showing PRDM14 expression in embryoid body (EB) sections throughout hPGCLC specification from PRDM14-AID-Venus fusion reporter cell line. hPGCLCs were induced by cytokines and EBs were collected at 12hrs, and on D1-D5. Representative images for 12hrs, D1, D2 and D4 are shown. hPGCLCs were marked by SOX17 and highlighted by a dashed line. Arrows show examples of SOX17+ cells that are PRDM14-negative at 12hrs and PRDM14-positive on D1 of differentiation. (**B**) Quantification of results from (A) including the D1-D5 time points. Data show the percentage of PRDM14+Venus+SOX17+ hPGCLCs as mean+/-SD of n=5 EB sections from one hPGCLC induction.

We also assessed PRDM14 expression in human gonadal PGCs, and unlike some previous reports^3,5^, we detected PRDM14 not only in the cytoplasm, but also in the nucleus of hPGCs (Suppl. Fig.1; Suppl. Fig. 1A). Nuclear localisation was also validated using IF on FACS-purified AP^+^cKIT gonadal hPGCs (Suppl. Fig.1B).

Next, we tracked PRDM14 expression dynamics upon hPGCLC induction by time-course IF (Fig.2A-B, Suppl. Fig.2). PRDM14 was uniformly expressed in 4i hESCs (Fig.1E) that are competent for hPGCLC-fate^3^. However, 12 hours (hrs) from the start of hPGCLC induction by cytokines, the levels of PRDM14 declined significantly in the SOX17+ hPGCLCs (Fig. 2A). The remaining PRDM14-Venus+ cells at 12hrs were SOX2+, unlike the nascent hPGCLCs, and thus likely represented neighbouring pluripotent cells (Suppl. Fig.2). The repression of PRDM14 was however transient, since we observed specific re-expression of PRDM14 in ~20% of SOX17+ cells on day 1 (D1); the proportion of PRDM14+SOX17+ cells continued to increase progressively, reaching ~40% on D2, ~80% on D3, ~90% on D4, and ~95% on D5 (Fig.2B). Of note, high SOX2 and PRDM14 levels persisted in the control cell aggregates in the absence of hPGCLC-inducing cytokines (Suppl. Fig.2). The transient loss followed by re-expression of PRDM14 in cells undergoing hPGCLC specification suggests that PRDM14 might have a vital role in this cell fate decision. We pursued this possibility by using inducible degrons for an acute destabilisation of the PRDM14 protein.

### Inducible degrons allow fast and reversible PRDM14 protein depletion

Most inducible loss-of-function approaches act at the transcriptional or post-transcriptional levels and often suffer from slow and incomplete protein removal^26–29^. Since PRDM14 is one of the core pluripotency factors in hESCs^13^, and exhibits dynamic changes during hPGCLC induction, conventional knockout approaches are unsuitable to study its functions in hPGCLCs. As PRDM14 expression commences within 24hrs of hPGCLC induction, we decided to use conditional degrons^30^ to achieve fast, inducible and reversible PRDM14 depletion at the protein level.

Auxin and jasmonate-inducible degrons (AID and JAZ, respectively), are plant pathways for ligand-induced targeted protein degradation^23, 24^ To harness AID and JAZ degrons in hESCs and hPGCLCs, we used CRISPR/Cas9 to tag the C-terminus of the endogenous PRDM14 with a 44-amino-acid AID^31, 32^, or a 43-amino-acid JAZ, fused to Venus (Fig.3A). Next, we employed the PiggyBac transposon system^33^ to deliver codon-optimised transgenes encoding cognate hormone receptors from rice (*Oryza sativa):* TIR1 for AID and COI1B for JAZ, allowing target degradation upon administration of auxin (indol-acetic acid, IAA) or coronatine (Cor), respectively^24, 32^.

**Figure 3.**
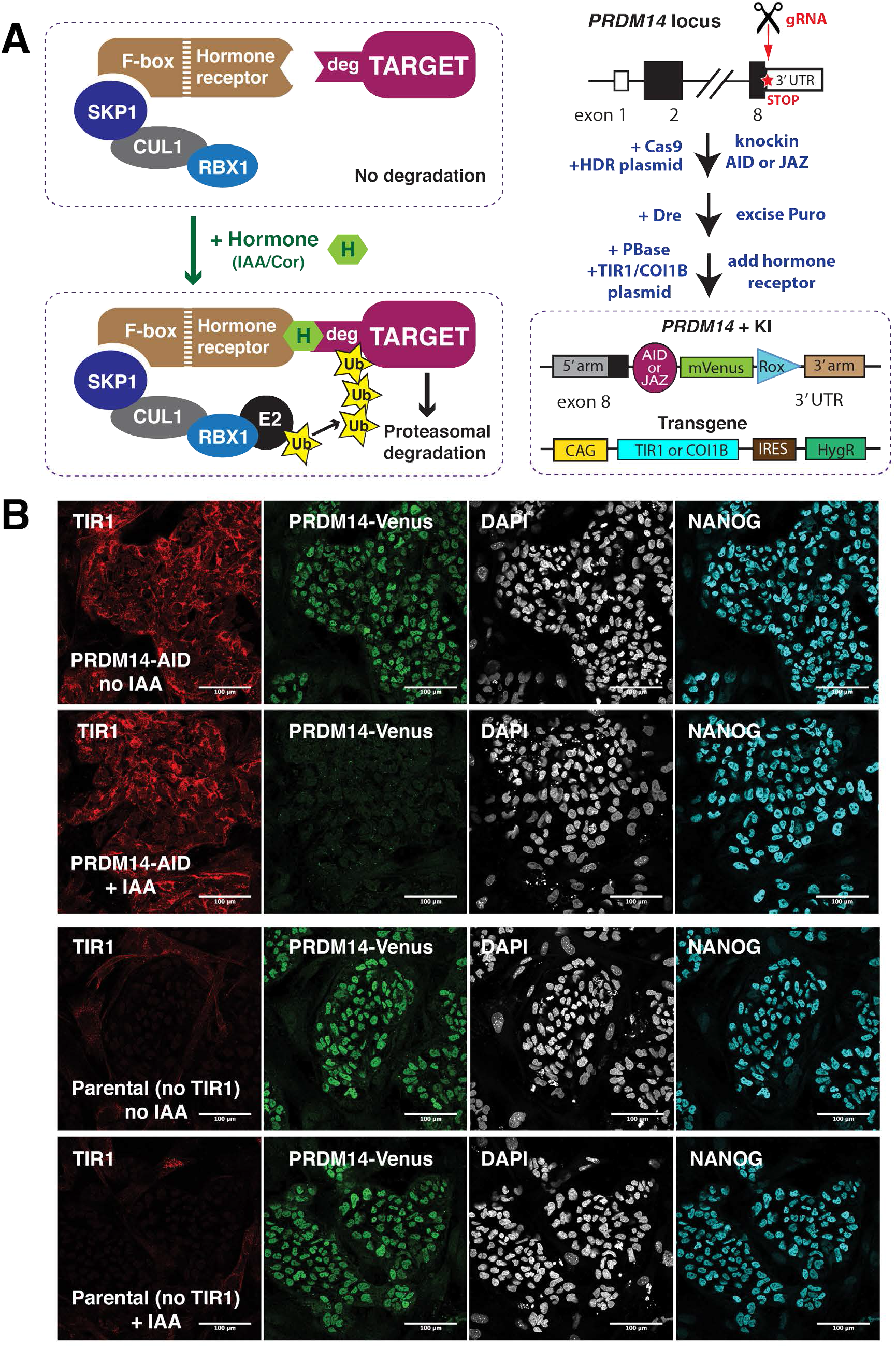
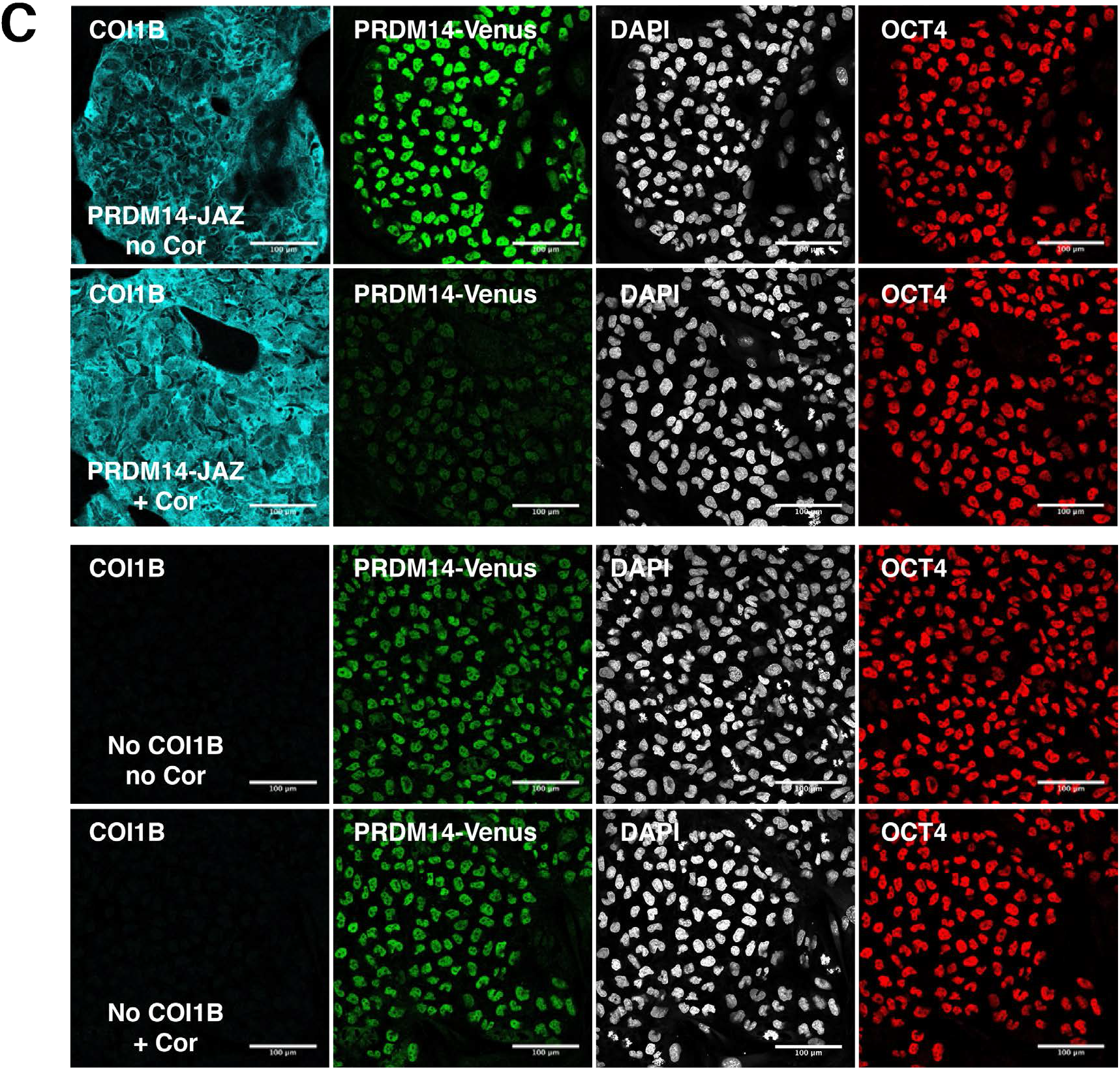
Auxin- and jasmonate-inducible degrons allow fast and homogenous PRDM14 protein depletion. (**A**) Scheme of AID and JAZ systems and the derivation of PRDM14-AID-Venus and PRDM14-JAZ-Venus cell lines. (**B**) IF showing PRDM14, NANOG and TIR1 (myc-tagged) expression and PRDM14-AID-Venus depletion upon 25 minutes of IAA treatment. Auxin- and jasmonate-inducible degrons allow fast and homogenous PRDM14 protein depletion. (**C**) IF showing PRDM14-Venus, OCT4 and COI1B (HA-tagged) expression and PRDM14-JAZ-Venus depletion upon 2 days of Cor treatment.

First, we tested AID- or JAZ-mediated PRDM14 depletion in hESCs by growing them with or without corresponding hormones and performing IF (Fig.3B-C, Suppl. Fig.3). PRDM14 and Venus fully colocalized and were both reduced upon the addition of IAA or Cor in hormone-sensitive, but not in parental lines (Fig.3B-C, Suppl. Fig.3A-D). Crucially, the AID system allowed reduction of Venus fluorescence to negligible levels, comparable to cells lacking the Venus knock-in (Suppl. Fig.3C). Time-course IF in hESCs revealed a rapid onset of PRDM14-AID-Venus depletion within 10 minutes of IAA treatment, reaching homogeneity within 25 minutes (Fig.3B, Suppl. Fig.3E). In the case of the JAZ degron, however, residual PRDM14-JAZ-Venus signal was detectable even after 2 days of Cor treatment (Fig.3C, Suppl. Fig.3D). Both systems are nevertheless reversible^24, 32^, and IAA wash-off restored PRDM14 levels in less than 2 hours (Suppl. Fig.3F). Altogether, inducible degrons allow fast and reversible PRDM14 depletion, with AID being superior in terms of speed and efficiency.

### Inducible PRDM14 degradation reduces hPGCLC specification

Next, we addressed the importance of PRDM14 for hPGCLC specification by adding IAA or Cor at the onset of hPGCLC differentiation (D0), followed by measuring the induction efficiency on D4 by recording the percentage of NANOS3-tdTomato^+^AP^+^ cells (Fig.4A). Notably, PRDM14 depletion using both approaches resulted in a significant reduction of hPGCLC induction efficiencies (by 70% and 30-60%, respectively) (Fig.4B-C, Suppl. Fig.4A-C). The effect was more pronounced in the case of the AID system, presumably due to its higher efficiency and faster kinetics of PRDM14 depletion. Therefore, we focused on the AID system to study PRDM14 further. Crucially, the observed phenotype was replicated using a PRDM14-AID in another hESC line (Suppl. Fig.4D).

**Figure 4.**
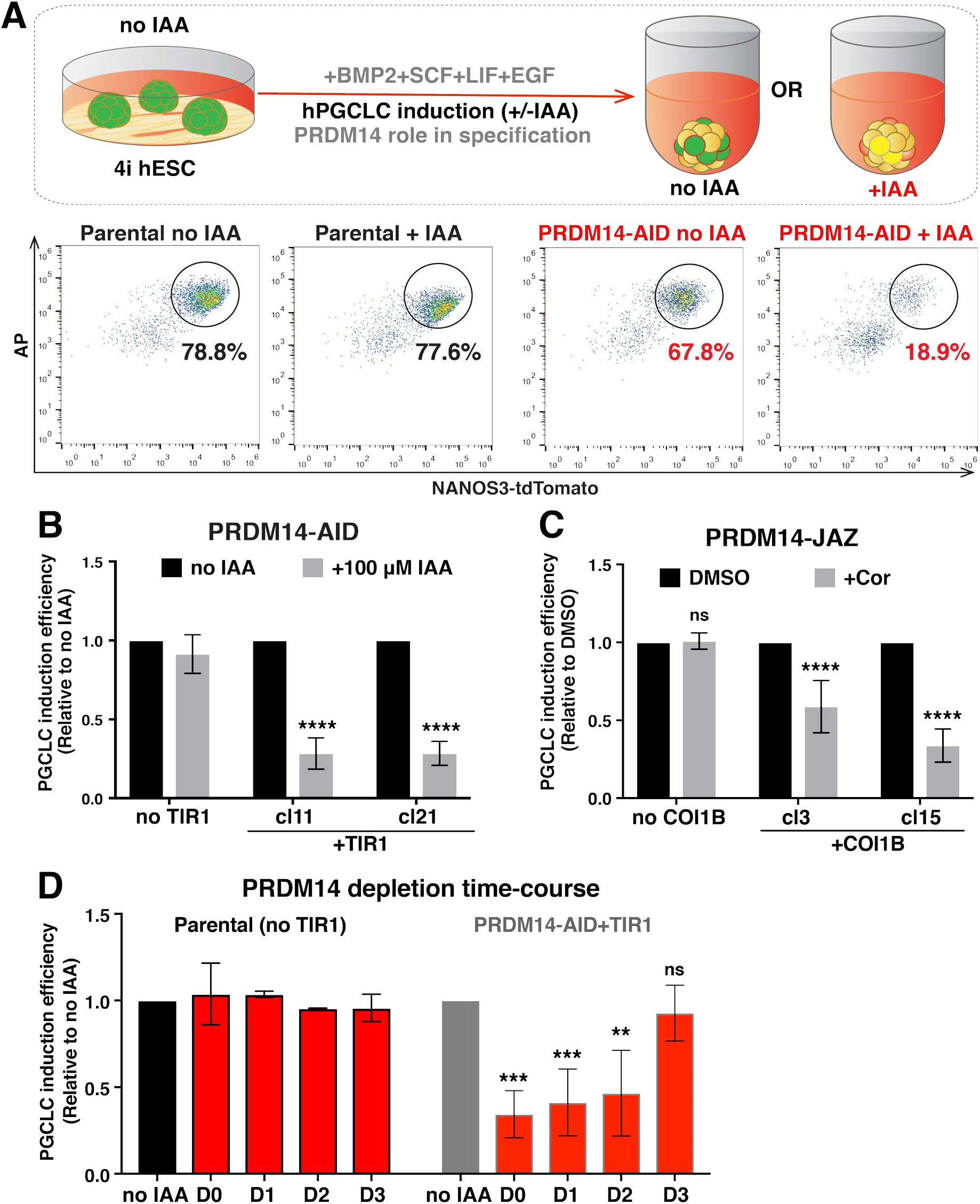
PRDM14 depletion using AID or JAZ degrons reduces hPGCLC specification efficiency. (**A**) Experimental design and representative flow cytometry plots showing reduced hPGCLC induction in hormone-sensitive clones, but not in the parental control. hPGCLCs were defined as NANOS3-tdTomato+AP+ cells. (**B**) Quantification of flow cytometry results for PRDM14-AID-Venus hPGCLCs induced with or without IAA (100μM). The induction efficiency of untreated cells (“no IAA”) was set to 1. Data show mean+/-SD for n=5-6 independent experiments per clone, ns – not significant, **** P<0.0001 (Two-way ANOVA followed by Sidak’s multiple comparison test). (**C**) Quantification of flow cytometry results for PRDM14-JAZ-Venus hPGCLCs. The induction efficiency of untreated cells (“no Cor”) was set to 1. Data show mean+/-SD for n=4 independent experiments per clone, ns – not significant, **** P<0.0001 (Two-way ANOVA followed by Sidak’s multiple comparison test). (**D**) Time-course of PRDM14 depletion. IAA (500 μM) was added on indicated days of hPGCLC induction from PRDM14-AID-Venus clones. Note that IAA was not washed off until the end of differentiation (D4), when hPGCLC induction efficiency was assessed by flow cytometry. Data show mean+/-SD of n=3 (2 independent experiments using 2 clones), or n=2 for “no TIR1” control, ns – not significant, ** P<0.01, *** P<0.001 (two-way ANOVA followed by Sidak’s multiple comparison test).

To further test AID efficacy in this context, we induced degradation of SOX17, a known hPGC and definitive endoderm (DE) regulator^3, 20, 34^. IAA addition fully abrogated hPGCLC and DE specification (Suppl. Fig.4E-G), confirming the key roles of SOX17 in these lineages. Notably, as a control, PRDM14 depletion had no adverse effect on the induction of definitive endoderm (DE) (data not shown), confirming the specificity of AID for our investigation.

To establish the time when PRDM14 is essential for hPGCLC specification, we performed a time-course of IAA supplementation. We performed depletion of PRDM14 by adding IAA starting on D0, D1 or D2, followed by analysis on D4. We noted a strong phenotype with reduction in hPGCLC specification (Fig.4D). By contrast, the addition of IAA on D3 had no detectable effect on the number of NANOS3-tdTomato^+^AP^+^ cells (Fig.4D). This suggests that either PRDM14 is dispensable after D2 or that its depletion at later time-points does not affect hPGCLC numbers, although transcriptional or epigenetic consequences cannot be excluded. Since PRDM14 starts being expressed in many hPGCLCs on D2 (Fig.2) and its loss from D2 to D4 yielded similar results to D0-D4 depletion (Fig.4D), it is likely that PRDM14 is most critical around D2 of hPGCLC specification. Therefore, PRDM14 might act to consolidate the germ cell network established by SOX17 and BLIMP1 from D1 of hPGCLC induction^3, 5^.

PRDM14 is highly expressed in human PGC-competent pluripotent cells, unlike in mice, where it is repressed at the equivalent stage^3, 35^. Notably, however, PRDM14 depletion in 4i hESCs for 1 passage prior to hPGCLC induction (Suppl. Fig.4H) had no effect on hPGCLC specification (Suppl. Fig.4I), indicating no detectable involvement of PRDM14 in the maintenance of competence for hPGCLC fate. To determine if PRDM14 is required for the acquisition of competence, we induced competence for hPGCLCs in hESCs via pre-mesendoderm (preME), which transiently displays hPGCLC potential^5^. However, when IAA was added to the preME medium and washed off before hPGCLC induction (Suppl. Fig.4H), we observed no effect on hPGCLC specification efficiency (Suppl. Fig.4J). Altogether, these data demonstrate that PRDM14 is important for hPGCLC specification but is dispensable for the acquisition and maintenance of the hPGCLC-competent state.

### Ectopic PRDM14 or desensitization to auxin rescues hPGCLC differentiation

To confirm the specificity of the observed phenotype, we attempted to rescue the endogenous PRDM14 depletion with ectopic *PRDM14,* using the ProteoTuner system, whose kinetics is comparable to that of AID^36^. PRDM14 transgene was fused to a destabilisation domain (DD) and thus continuously degraded by the proteasome but stabilised by Shield-1 ligand^36, 37^. Interestingly, overexpression of PRDM14 on D0, but not on D1 could fully rescue hPGCLC specification efficiency and even enhanced hPGCLC specification compared to “no IAA” control (Fig.5A). This suggests that the transient PRDM14 repression observed at the onset of hPGCLC specification (Fig.2B) is not prerequisite for germ cell induction.

**Figure 5.**
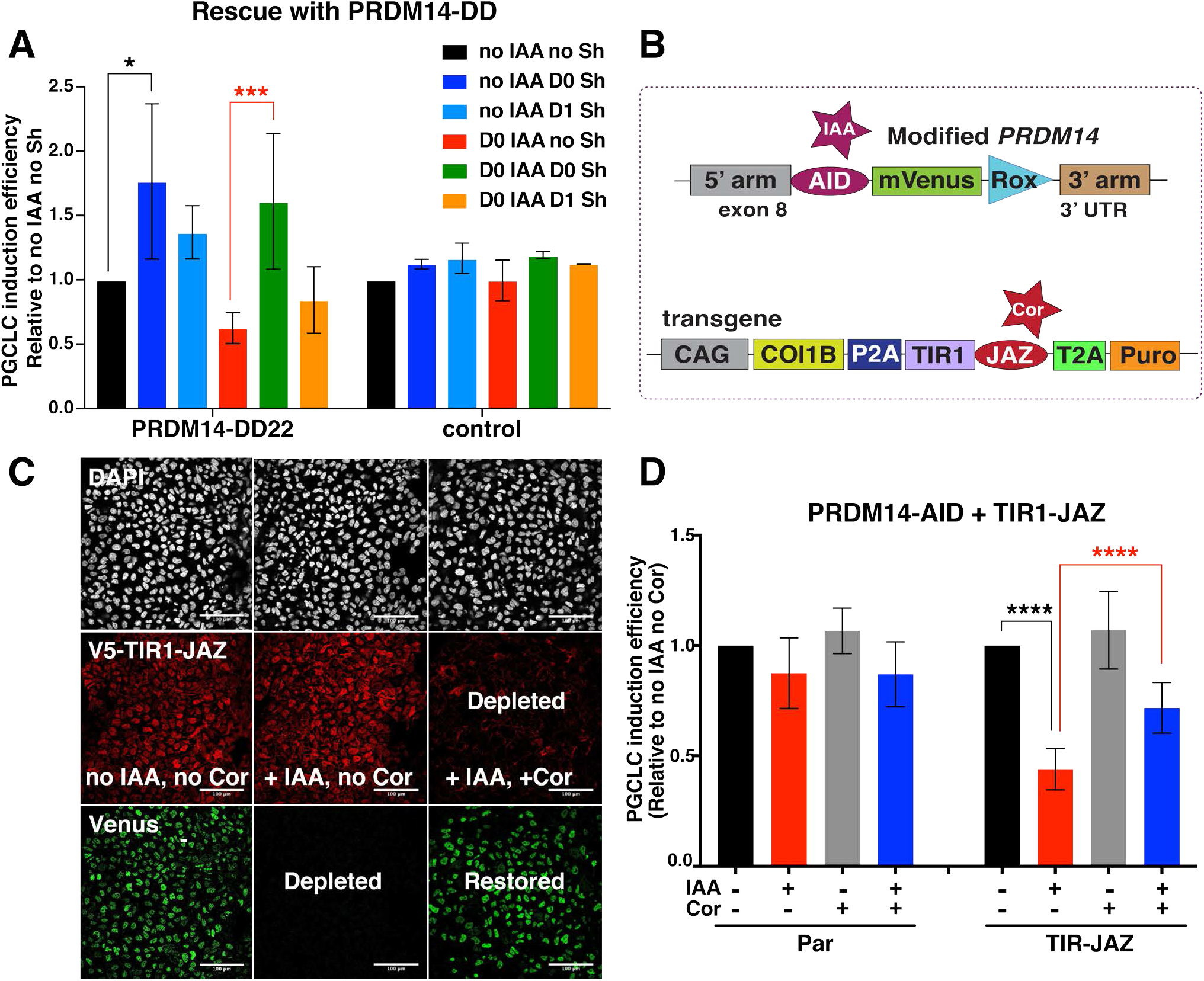
Ectopic PRDM14 expression or desensitisation to auxin through TIR1 degradation rescue hPGCLC specification phenotype. (**A**) IAA and/or Shield-1 (Sh) were added on indicated days of differentiation to deplete the endogenous PRDM14 or stabilise the PRDM14-DD transgene, respectively. “Control” cell line lacks both the AID knock-in and the PRDM14-DD transgene. Data show mean+/-SD for n=4 (PRDM14-DD22) or n=2 (control) independent experiments, * P<0.05, ** P<0.01 (Two-way ANOVA followed by Sidak’s multiple comparison test separately within “no IAA” or “D0 IAA” groups). (**B**) Scheme of the double AID/JAZ degron design. Endogenous PRDM14 is under the control of the AID degron as above, while the stability of TIR1 hormone receptor is controllable by the JAZ degron. (**C**) AID/JAZ validation by IF in hESCs. Simultaneous IAA and Cor supplementation depletes TIR1 and restores PRDM14, as shown by IF staining for PRDM14-AID-Venus and TIR1(V5-tagged). Nuclei were counterstained by DAPI. Scale bar is 100 μm. (**D**) Desensitisation to auxin via the AID/JAZ degron switch rescues hPGCLC induction efficiency. Data show mean+/-SD for n=4 (no TIR1 control) or n=11 (TIR-JAZ, 3 clones combined), **** P<0.0001 (Two-way ANOVA followed by Sidak’s multiple comparison test).

We then asked if we could restore the endogenous PRDM14 levels by blocking auxin perception. To achieve this, we designed an AID-JAZ degron switch, where PRDM14-AID-Venus is degraded by IAA supplementation, while TIR1 is fused to JAZ and thus depleted by Cor (Fig.5B). IF analysis in hESCs confirmed that simultaneous administration of both IAA and Cor desensitised hESCs to IAA via TIR1 degradation and replenished the PRDM14-Venus pool (Fig.5C). In line with previous experiments, these cell lines also showed decreased hPGCLC specification efficiency upon IAA treatment (Fig.5D). Crucially, in the presence of both IAA and Cor, hPGCLC differentiation efficiency was significantly restored (Fig.5D). This confirms the causative role of PRDM14 depletion in hPGCLC specification phenotype and exemplifies the use of orthogonal degron approaches for rescue design.

### PRDM14-defìcient hPGCLCs show aberrant transcriptome

Next, we examined the transcriptome of PRDM14-deficient hPGCLCs by performing RNA-sequencing (RNA-seq) on NANOS3-tdTomato+AP+ hPGCLCs differentiated with or without IAA for 4 days from 2 hormone-sensitive clones and the parental control lacking TIR1. We also addressed the role of PRDM14 in pluripotency regulation, by analysing the transcriptome of AP+ competent hESCs grown with or without IAA for 1 passage (3 days). Importantly, the parental (“no TIR1”) control showed no differentially expressed genes (DEGs) in IAA-treated hESCs, and only 2 upregulated genes in hPGCLCs (*CYP1B1* and *LRAT*), indicating minimal non-specific effects of auxin on the target cells’ transcriptome (Suppl. Table 9). RNA-seq on PRDM14-depleted hESCs identified 106 upregulated and 64 downregulated genes (Fig.6A and Suppl. Table 9). Crucially, our findings concur with a previous study on PRDM14 in different hESC lines and culture conditions^13^ (Fig.6C), with downregulation of pluripotency genes (e.g. *OCT4, NANOG, TDGF1,* and *GAL),* and upregulation of differentiation markers, particularly of the neural lineage (e.g. *MAP2, MAP6* and *SOX2*) (Fig.6A, Suppl. Fig.5E, Suppl. Table 9). Gene ontology (GO) analysis reflected reduction in selfrenewal (enriched downregulated term “somatic stem cell maintenance”) and upregulation of prodifferentiation genes (enriched terms “nervous system development” and “embryonic pattern specification”) (Fig.6F).

**Figure 6.**
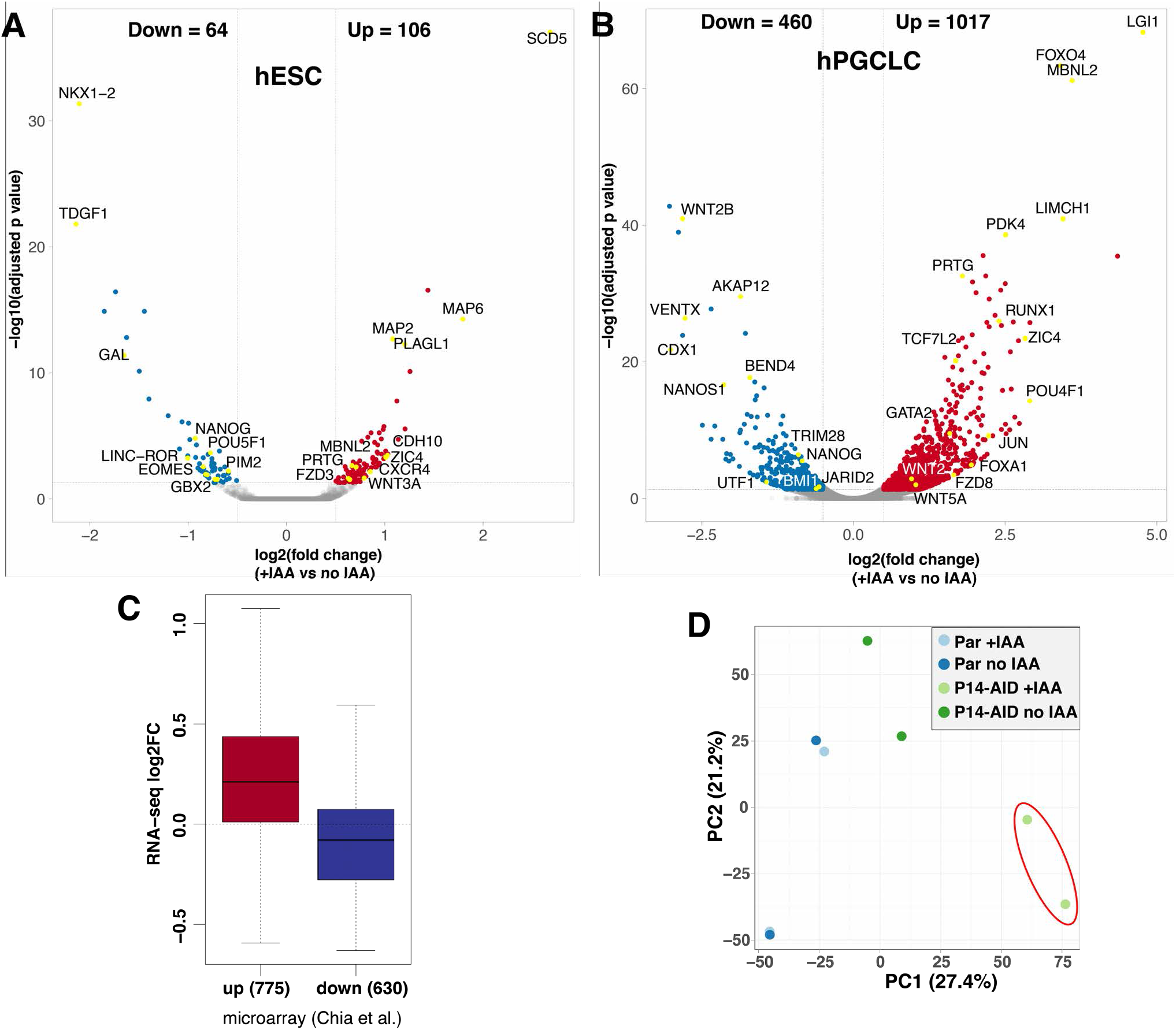
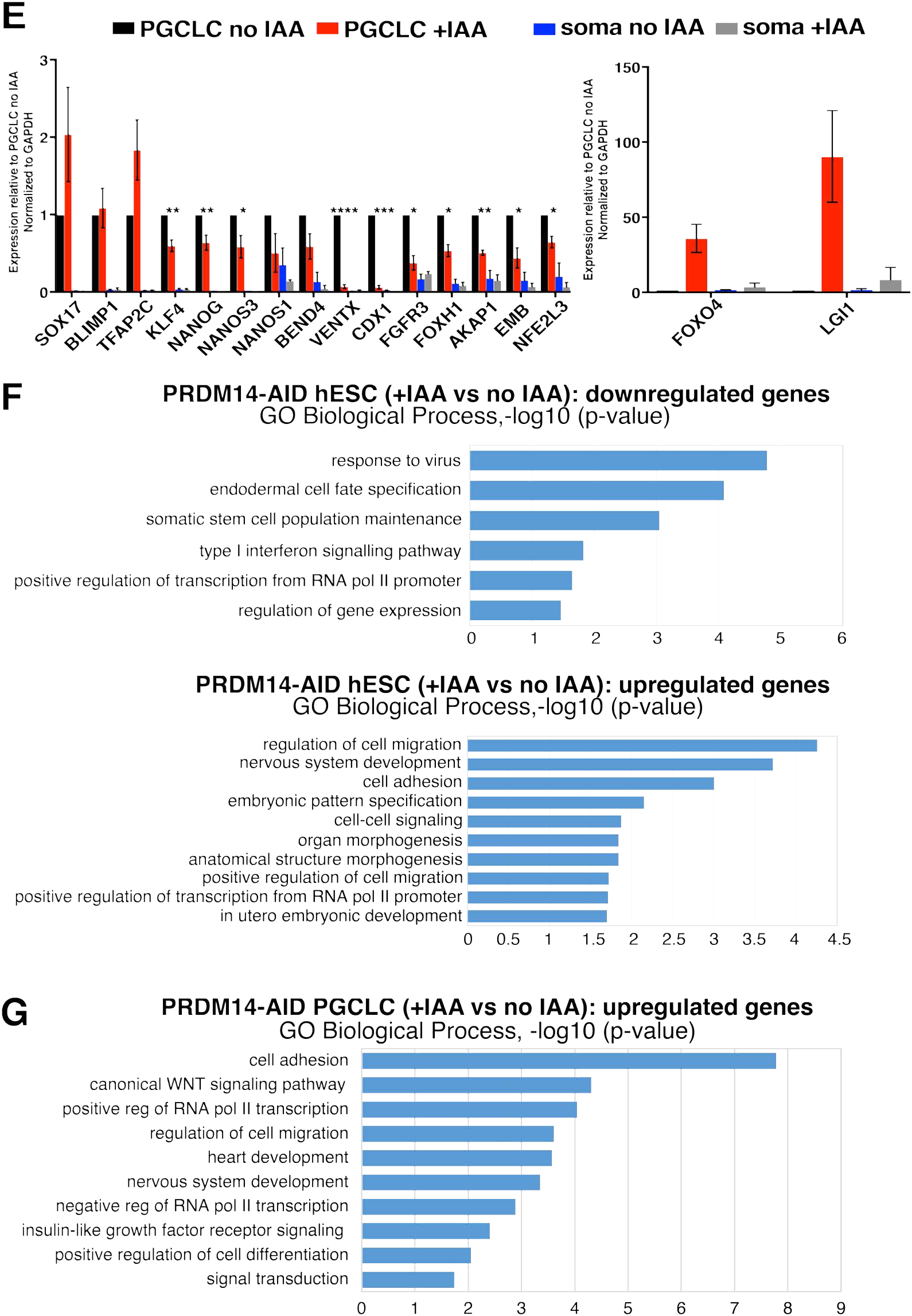
RNA-seq reveals transcriptional changes in PRDM14-deficient hESCs and hPGCLCs. (**A**) Volcano plot showing down- (log2FC<(−0.5), padj<0.05) and upregulated (log2FC>0.5, padj<0.05) genes in hPGC-competent hESCs upon PRDM14 loss. PRDM14-AID hESCs were treated with IAA for 1 passage (3 days) and sorted as AP+ cells for RNA-seq along with “no IAA” control. (**B**) Volcano plot showing down- (log2FC<(−0.5), padj<0.05) and upregulated (log2FC>0.5, padj<0.05) genes in hPGCLCs upon PRDM14 loss. PRDM14-AID hPGCLCs were specified with or without IAA for 4 days and sorted as NANOS3tdTomato^+^AP^+^ cells for RNA-seq along with “no IAA” control. (**C**) Boxplot overlaying down- (log2FC<(−0.5), padj<0.05) and upregulated (log2FC>0.5, padj<0.05) genes upon *PRDM14* knockdown in conventional hESCs (microarray data from^13^) and RNA-seq analysis in PRDM14-AID-depleted PGC-competent (4i) hESCs from this study. Note that the majority of differentially expressed genes show consistent changes. (**D**) Principal component analysis (PCA) of hPGCLC RNA-seq samples. pRDM14-depleted hPGCLCs (cl11 and cl21; highlighted in red) separate from the other samples along principal component 1 (PC1). RNA-seq reveals transcriptional changes in PRDM14-deficient hESCs and hPGCLCs. (**E**) qPCR validation of selected DEGs from RNA-seq. hPGCLCs from cl11 and cl21 were sorted as NANOS3-tdTomato^+^AP^+^ cells, while soma denotes the double-negative population from the same experiments. Strongly upregulated genes are plotted separately. Data show gene expression relative to no IAA hPGCLCs and normalized to *GAPDH,* mean+/-SEM of n=2-5 independent experiments, * P<0.05, ** P<0.01, *** P<0.001, **** P<0.0001 (multiple t-tests). The parental (“no TIR1”) control is shown in Suppl. Fig.5D. (**F**) Gene ontology (GO) analysis on down- and upregulated genes in PRDM14-deficient hESCs. Top non-redundant GO terms are shown as –log10(p-value). (**G**) GO analysis on upregulated genes in PRDM14-deficient hPGCLCs. Top non-redundant GO terms are shown as –log10(p-value). For complete list of GO terms see Supplementary Table 10.

Notably, there was a stronger response to PRDM14 loss in hPGCLCs, with 1017 upregulated and 460 downregulated genes in “+IAA” vs “no IAA” samples (Fig.6B), and only a small overlap between the affected genes in hESCs and hPGCLCs (Suppl. Table 9). Principal component analysis (PCA) and unsupervised hierarchical clustering of hPGCLC transcriptome confirmed that parental control, irrespective of IAA treatment, clustered with untreated hormone-sensitive clones, away from the PRDM14-depleted hPGCLCs (Fig.6D and Suppl. Fig.5A). Crucially, PRDM14-deficient hPGCLCs showed downregulation of many key hPGC- and pluripotency-related genes, including *UTF1, NANOG, NANOS1, LIN28A,* and *TRIM28.* Indeed, gene set enrichment analysis (GSEA) against 123 core PGC genes defined previously^5^, showed lower enrichment score of mutant hPGCLCs compared to the control (Suppl. Fig.5B). The levels of key hPGC regulators, *SOX17* and *PRDM1,* however did not change (Fig.6E and Suppl. Fig.5D), which is consistent with their upregulation prior to *PRDM14* in nascent hPGCLCs (Suppl. Fig.2 and^3^). Ectopic *PRDM14* could partially revert the transcriptional changes in mutant hPGCLCs (Suppl. Fig.5C), in line with rescued hPGCLC specification phenotype (Fig.5A).

GO analysis on genes derepressed in PRDM14-deficient hPGCLCs identified enrichment of terms related to WNT signalling, as well as heart and nervous system development (Fig.6G). A similar response occurs in hPGCLCs specified without *BLIMP1* or *TFAP2C* (Kojima et al., 2017; Tang et al., 2015). Indeed, pairwise comparisons revealed a significant overlap between both up- and downregulated targets, suggesting potential combinatorial roles of PRDM14 with BLIMP1 and TFAP2C. Accordingly, 115 genes were downregulated in all three mutants (including germ cell markers *UTF1, NANOG, AKAP12, NANOS1,* and *TRIM28),* while 281 shared upregulated genes were mostly implicated in morphogenesis, WNT signalling, as well as cell migration and adhesion (Suppl. Fig.5F-G and Suppl. Table 11).

### PRDM14 regulates target gene expression mainly through promoter binding

To identify direct targets of PRDM14 in hPGCLCs and hESCs, we performed chromatin immunoprecipitation with high-throughput sequencing (ChIP-seq). We took advantage of the PRDM14-AID-Venus line to use a ChIP-grade anti-GFP antibody for PRDM14-Venus immunoprecipitation, which proved more efficient than anti-PRDM14 antibodies (Suppl. Fig.6A). Importantly, auxin treatment led to a significant loss of signal (Suppl. Fig.6A), which allowed us to use ChIP-seq on IAA-treated hESCs as a control.

Overall, ChIP-seq identified 6486 consensus PRDM14 peaks: 4206 in hPGCLCs and 3319 in hESCs (Suppl. Table 12). *k*-means clustering partitioned the dataset into 5 distinct clusters (Fig.7A), separating peaks specific to hESCs (742 peaks, cluster 4) or hPGCLCs (1331 peaks, cluster 5) or shared between the two (clusters 1-3). Note that the strongest ChIP-seq enrichment persisted in IAA-treated hESCs (clusters 1-2) with 1728/3319 hESC binding sites detectable in hESC+IAA, albeit with significantly lower signal (Fig.7A and Suppl. Table 12). Such persistent regions might be more tightly bound, and/or require longer auxin exposure to eliminate PRDM14 binding completely.

Notably, PRDM14 in both hESCs and hPGCLCs predominantly binds within 1 kilobase (kb) of transcription start sites (TSS) (Suppl. Table 12), with ~30% of peaks spanning annotated promoters (Fig.7B), which agrees with a previous ChIP-seq in hESCs^13^. By contrast, PRDM14 binds predominantly to distal genomic regions in mice, and only 4-10% of peaks are located within 1 kb from the TSS in mESCs^16, 38^.

Motif analysis^39^ confirmed that the conserved PRDM14 motif was top-ranking in both hESCs and hPGCLCs (Fig.7C). Notably, TFAP2C motif was the second most enriched within hPGCLC-specific targets, suggesting cooperation between the two factors (Fig.7C), consistent with the overlapping targets from RNA-seq (Suppl. Fig.5F). We also found significant enrichment of BLIMP1 and SOX motifs in hPGCLCs, whereas unique hESC peaks were enriched in the OCT4-SOX2 motif (Fig.7C). Altogether, this indicates that PRDM14 co-occupies genomic targets with core TFs specific to hPGCs or pluripotent cells, respectively.

Next, we correlated ChIP-seq peaks mapping within 100 kb of the TSS with the corresponding changes in gene expression from RNA-seq (Fig.7D, Suppl. Fig.5H and Suppl. Table 12). In hPGCLCs, 314 peaks were associated with 168 (17%) upregulated genes and 93 peaks with 47 (10%) downregulated genes (Suppl. Table 12). GO analysis confirmed that genes related to WNT signalling, heart and brain development are among the direct PRDM14 targets (Suppl. Fig.6B). The hESC-specific targets included *SOX2, ERBB3* and *GBX2,* while hPGCLC-specific peaks were found near *SOX17, TFAP2C* and *NANOS3* among others (Fig.7E-F). Whereas PRDM14 binds to the regulatory elements of *SOX17,* PRDM14 depletion did not significantly affect SOX17 expression in hPGCLCs (Fig.6E and Suppl. Table 9) presumably because other factors sustain its expression.

### Distinct molecular functions of PRDM14 in mouse and human

Next, we asked if the molecular roles of PRDM14 are conserved between mouse and human, by comparing the protein-coding transcriptomes of PRDM14-AID hESCs and hPGCLC, with the equivalent mouse *Prdm14*^-/-^ cells^11^ (Suppl. Fig.7A-B, Suppl. Table 13). PGC-competent mouse epiblast-like cells (mEpiLCs) showed very few DEGs since *Prdm14* is repressed in these cells^11^. Only two genes (*CDH4* and *HS6ST2*) were derepressed in both mESCs and hESCs, and 3 genes (*DNMT3B, SPRY4* and *FHL1*) showed opposite changes, probably reflecting the broader differences between mESCs and hESCs^40^. Comparison of PRDM14-depleted hPGCLCs with D6 *Prdm14^-/-^* mPGCLC identified a limited subset of potentially conserved PRDM14 targets: 36 up- and 13 downregulated genes (Suppl. Fig.7A), but none have hitherto been implicated in germ cell biology. Furthermore, the expression of 39 DEGs was anti-correlated in hPGCLC and mPGCLCs (Suppl. Fig.7A).

In mPGCs, PRDM14 initiates global epigenetic reprogramming, in part through the repression of *Uhrf1* to allow passive DNA demethylation^11^; this gene is also repressed in hPGCs and hPGCLCs^41^. A small subset of PRDM14-depleted proliferating SOX17+ cells failed to downregulate UHRF1, potentially suggesting a conserved regulation (Suppl. Fig.7C).

PRDM14 alone is sufficient to induce mPGCLCs fate^8^; however, this was not the case for hPGCLC specification using either doxycycline-inducible or ProteoTuner systems (Suppl. Fig.7D), consistent with the observation that upregulation of SOX17 and TFAP2C precedes that of PRDM14 (Fig.2). Altogether PRDM14 evidently plays an important role shortly after the initiation of hPGCLC fate, but its function is distinct from that in mPGCs.

**Supplementary Figure 7.**
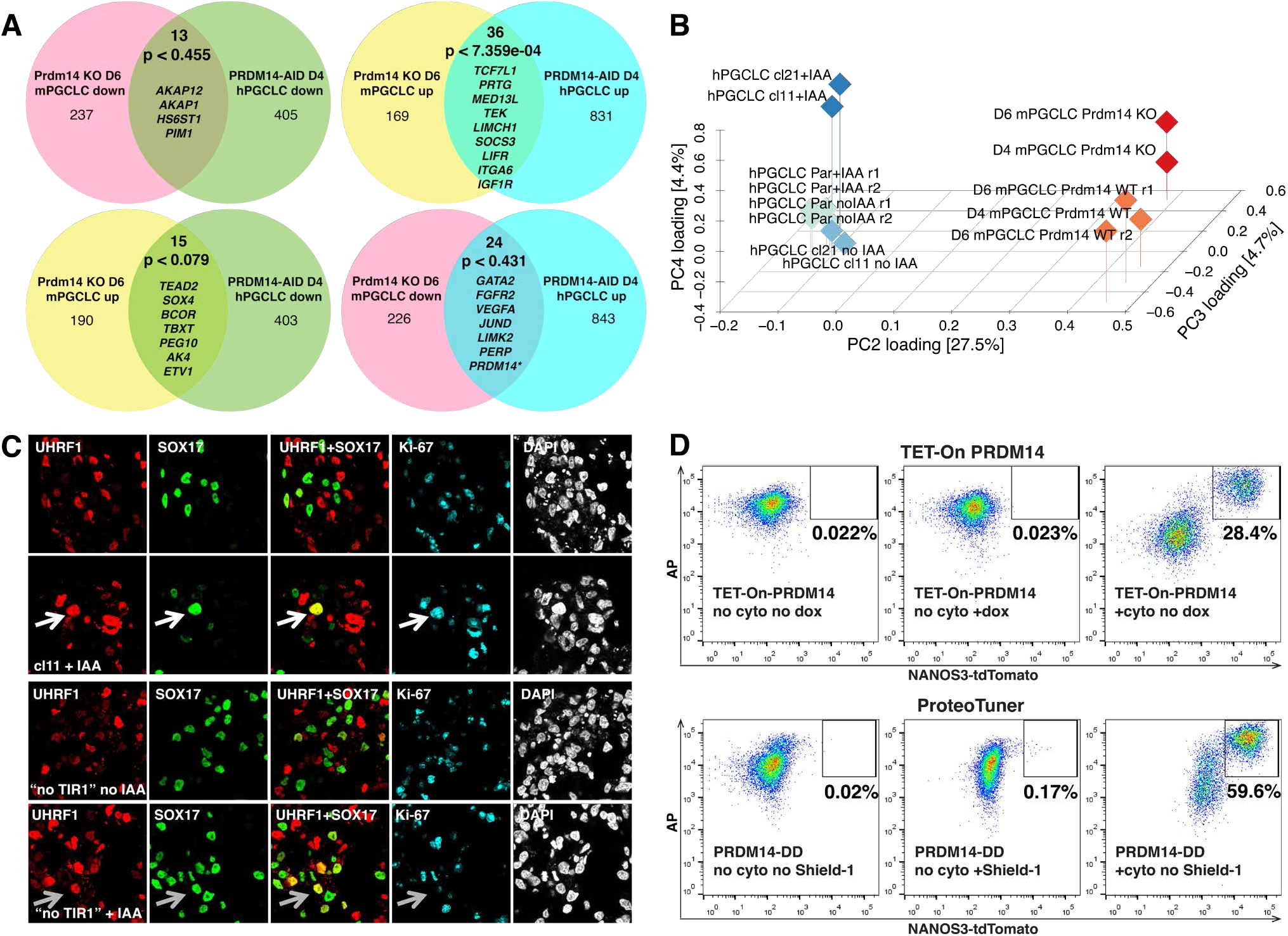
Comparison of molecular functions of PRDM14 in mouse and human hPGCLCs. (**A**) Venn diagrams comparing transcriptional changes in *Prdm14^-/-^* mouse PGCLCs and PRDM14-AID hPGCLCs (+IAA) relative to corresponding WT controls. P-values were obtained by pairwise hypergeometric tests. Examples of overlapping DEGs are shown. * In PRDM14-AID hPGCLCs PRDM14 is depleted only at the protein level, while the transcript is still present and even upregulated, presumably due to autoregulation. (**B**) 3D PCA plot showing distinct transcriptional response of mouse and human PGCLCs to the absence of PRDM14. Data from Suppl. Table 13 with logFC>1 were used. Note that mouse WT and KO samples separate by PC4, while human samples induced with or without PRDM14 separate along PC3. Par – parental control lacking TIR1; r1 and r2 – replicates 1 and 2. (**C**) IF analysis of UHRF1 expression in D4 PRDM14-AID EBs. Putative hPGCLCs are marked by SOX17, proliferating cells are marked by Ki-67. White arrows highlight an example mutant proliferating hPGCLC that retains UHRF1. Note that UHRF1+SOX17+ cells in the “no TIR1” control (grey arrows) are Ki67-negative. (**D**) Representative flow cytometry plots showing that ectopic PRDM14 induced by either dox or Shield-1 does not induce hPGCLC induction without BMP2 and other cytokines (“no cyto”). Dox or Shield-1 were added on D0 of differentiation.

## Discussion

Using two acute protein depletion strategies, combined with rescue, transcriptomic and ChIP-seq experiments, we demonstrate that PRDM14 is required for hPGCLC specification and represses somatic differentiation while promoting germline fate (Suppl. Fig.8). Strikingly, the molecular function of PRDM14 in the human germline has diverged significantly compared to mPGCs. Indeed, the sets of targets regulated by PRDM14 in the two species are vastly different (Suppl. Fig.7A-B) and, unlike in the mouse^8^, PRDM14 alone cannot induce hPGCLCs (Suppl. Fig.7D). The regulatory network for hPGC specification is altogether distinct from that in mice, with the recently established critical role of SOX17, and the notable repression of SOX2^3, 20^. Our study of PRDM14 provides further insights on the divergence of the molecular basis of germ cell fate determination in mouse and human.

PRDM14 is specifically upregulated in the nucleus of nascent hPGCLC from D1 of germ cell induction, following the expression of SOX17 and TFAP2C (Fig.2). PRDM14 depletion strongly reduces the efficiency of germ cell specification (Fig.4) and results in an aberrant transcriptome of the PRDM14-deficient hPGCLCs (Fig.6). The effects of PRDM14 depletion during hPGCLC specification, resemble those observed upon the loss of *TFAP2C* or *BLIMP1* (Suppl. Fig.5F). Furthermore, ChIP-seq analysis revealed high enrichment of TFAP2C and BLIMP1 motifs within hPGCLC-specific PRDM14 targets (Fig.7C), which suggests combinatorial roles of the three regulators during the induction of hPGC-specific genes and the repression of somatic markers. However, unlike in mPGCs^11, 42^, many prominent germ cell genes, such as *TFAP2C, PRDM1* and *DND1,* are not the targets of PRDM14 in hPGCLCs. Furthermore, while the PRDM14 motif is conserved, PRDM14 binds predominantly to gene promoters in hESCs and hPGCLCs (Fig.7B and^13, 16, 38^), but to distal regulatory elements in mice^16, 38^. The notable lack of overlap might be dictated by different protein partners or by the divergence of the protein itself, although there is significant functional conservation of the protein, and human PRDM14 rescues the lack of its mouse orthologue in mESCs^15^.

The context-dependent roles of PRDM14 are also exemplified by distinct set of target genes in hESCs, where it cooperates with OCT4 and SOX2 to sustain pluripotency, and limits neuronal differentiation (Suppl. Fig.8, Fig.6A, Suppl. Fig.5E and Fig.7C). This and other studies highlighted the divergent roles of PRDM14 in mouse and human pluripotent cells^13, 15, 16, 38^. Furthermore, there are significant human–mouse differences during early embryo development, at the time of PGC specification^43, 44^; altogether these differences likely contribute to the species-specific modes of PGC specification. Notably, we found an even more pronounced transcriptional phenotype upon PRDM14 loss in hPGCLCs than in hESCs (Fig.6). Note that hESCs were, however, in a ‘steady state’ of self-renewal, while hPGCLCs were undergoing cell fate transitions at the time of PRDM14 depletion. The persistence of PRDM14 at some loci even after 3 days of IAA treatment (Fig.7A), is reminiscent of a significant (27%) retention of CTCF-AID peaks that were reported after 2 days of exposure to IAA^32^. Since most other inducible loss-of-function approaches display lower efficiency and slower kinetics, it is likely that they may result in even slower protein depletion from chromatin.

The distinct roles of PRDM14 in mouse and human PGCs, further illustrate the impact of the species-specific regulatory networks for PGC fate. For example, expression of SOX17 and SOX2 is mutually exclusive in germ cells of human and mouse, respectively. Whereas SOX17 is essential for hPGC fate^3^, SOX2 promotes mPGC specification and survival^45^. By contrast, repression of SOX2 is apparently prerequisite for the initiation of hPGCLC fate^1^, since ectopic expression of SOX2 abrogates hPGCLC specification^46^. In the absence of BMP signalling, which initiates hPGCLC specification, high expression of SOX2 and PRDM14 is maintained (Suppl. Fig.2). It is possible that PRDM14 expression in the hPGCLC-competent cells is downstream of SOX2. The repression of SOX2 upon the initiation of hPGCLC in response to BMP signalling might therefore explain a transient loss of PRDM14. Indeed, in hESC where SOX2 binds to PRDM14 promoter^47^, loss of *SOX2* leads to *PRDM14* downregulation^48^. The observed re-expression of PRDM14 in hPGCLC follows after the upregulation of SOX17 and BLIMP1 at the onset of hPGCLC specification (Fig.2 and Suppl. Fig.2), which suggests a molecular shift in the regulation of PRDM14 expression in germ cells.

In mPGCs, PRDM14 promotes DNA demethylation, in part by repressing *Uhrf1,* and initiates the reduction of H3K9me2 through *Ehmt1* downregulation^9^. How the TF network might control the epigenetic resetting in hPGCs is less clear, but some essential features of epigenetic reprogramming are evident in hPGCs, including the repression of *UHRF1* and EHMT2^41^. PRDM14-deficient hPGCLCs indicated derepression of UHRF1 (Suppl. Fig.7C) but not of EHMT2 in a subset of mutant hPGCLC. Further studies are required to establish how precisely the epigenetic resetting is initiated in hPGCs and a potential role for PRDM14, if any, in this critical process.

Our study shows the importance of the use of rapid and comprehensive PRDM14 depletion for the phenotype unmasking. A previous study of PRDM14 in hPGCLCs used a partial knockdown at a later time-point (D2 BLIMP1+SOX17+ precursors), where the homogeneity and speed of PRDM14 depletion was not monitored^19^. We demonstrate the utility of AID and JAZ degrons to deplete endogenous proteins to study human cell fate decisions. Simultaneous use of both degrons allows independent control of two proteins, and the construction of degron switches as shown here (Fig.5B-D). The knock-in of AID/JAZ together with a fluorescent reporter facilitates protein expression analysis and allows the use of the anti-GFP (or other epitope tags) antibody for IF and ChIP. Furthermore, tissue-specific TIR1 expression can allow spatial control over protein stability, as shown in *Caenorhabditis elegans* and *Drosophila melanogaster* without detectable side effects^49, 50^.

Neither mouse nor human adult tissues express PRDM14 except in some types of cancer^14, 51^, indicating the importance of studying the role of PRDM14 in normal embryogenesis. Inducible degrons offer a more precise control over proteins when studying the roles of critical TFs undergoing dynamic changes during normal and malignant development.

## Acknowledgements

We thank Dr. Elphège Nora for kindly providing reagents (plasmids containing AID degron and TIR1 sequences) and guidance for the establishment of the auxin-inducible degron. We are grateful to Dr. Ran Brosh for sharing the reagents (plasmids containing JAZ degron and COI1B sequences) and advice on the jasmonate-inducible degron. We thank Dr. Richard Butler for writing the Fiji plugin for fluorescence intensity quantification. We gratefully acknowledge Dr. Toshihiro Kobayashi, Dr. Naoko Irie and Dr. Ufuk Günesdogan for their useful advice throughout the project. AS was funded by the 4-year Wellcome Trust PhD Scholarship, MAS holds a Wellcome Senior Investigator Award, and grant from the MRC. The Gurdon Institute is supported through core funding from the Wellcome Trust and Cancer Research UK. WHG was supported by an EMBO long term fellowship (ALTF 263-2014). WWCT holds a Croucher Fellowship for Postdoctoral Research.

## Author contributions

The study was conceived and designed by AS and MAS. AS performed most experiments; WWCT and SD performed most bioinformatics analyses; WWCT performed IF on human gonadal sections. WHG helped with ChIP experiments. The study was supervised by MAS. The manuscript was written by AS and MAS with contributions from most authors.

## Competing interests

The authors declare no competing interests.

## Materials and Methods

### Human embryonic stem cell culture

PGC-competent 4i hESCs (H9^1^, WIS2-NANOS3-T2A-tdTomato (N3)^2^ and cell lines derived from them) were cultured as in^3^ on irradiated mouse embryonic fibroblasts (MEFs) (GlobalStem) in 4i medium (Suppl. Table 1). Media were replaced every day. hESCs were passaged by single-cell dissociation using 0.25% Trypsin-EDTA (GIBCO). 10 μM ROCK inhibitor (Y-27632, TOCRIS) was added for 24 hours after passaging.

Conventional hESCs (used for hPGCLC differentiation via pre-mesendoderm) were grown in Essential 8 (E8)^4^ in plates pre-coated with 5 μg/ml vitronectin for at least 1 hour. Media were replaced every day. Cells were passaged in clumps using 0.5 mM EDTA in PBS. All reagents for E8 hESC culture were from Thermo Fisher Scientific.

For transgene inductions in hESCs or during differentiation, 1 μg/ml doxycycline (dox, Sigma) and/or 0.5 μM Shield-1 (Clontech) were added to media, where specified. For depletion of AID-fused proteins, auxin (indole-3-acetic acid sodium salt, IAA, Sigma) was used at 100 μM (in H2O) unless otherwise specified. For depletion of JAZ-fused proteins, coronatine (Cor, Sigma) was used at 50 μM (in DMSO) in all experiments. DMSO was used as vehicle control for experiments with Cor.

### PGC-like cell (PGCLC) and definitive endoderm (DE) induction

To induce hPGCLCs^3^, 4i hESCs were trypsinized, filtered and plated into ultra-low cell attachment U-bottom 96-well plates (Corning, 7007) at 4,000 cells/well density in 100 μl PGCLC medium (Suppl. Table 2)^2^. For induction of large quantities of hPGCLCs (for ChIP-seq and qPCR experiments), 6-well EZSPHERE microplates (ReproCELL) were used (500,000 cells/well in 3 mL PGCLC medium). The plates were centrifuged at 300g for 3 minutes and placed into a 37 °C 5% CO2 incubator until embryoid body (EB) collection for downstream analysis. Reporter fluorescence intensities were monitored daily throughout differentiation using Olympus IX71 microscope.

For hPGCLC induction from E8 (conventional) hESCs^2^, pre-mesendoderm (preME) competent state was induced by seeding 200,000 trypsinised single cells per well of a vitronectin-coated 12-well plate and culturing for 12 hours in mesendoderm (ME) medium (Suppl. Table 3). PreME cells were then trypsinised, filtred and induced similar to 4i hESCs, as described above.

For definitive endoderm (DE) induction^2^, mesendoderm (ME) was first obtained from E8 cells by 36-hour exposure to ME medium (Suppl. Table 3); ME was then washed with PBS followed by 48-hour culture in the DE medium (Suppl. Table 4).

### Flow cytometry and fluorescence-activated cell sorting (FACS)

Flow cytometry and FACS were performed as in^3^. At least 6 EBs were washed in PBS and dissociated with 0.25% Trypsin-EDTA for 10 min at 37°C. Cells were washed, resuspended in FACS buffer (3% FBS in PBS) and incubated with anti-AP and anti-CD38 antibodies specified in Suppl. Table 5 for 30-60 minutes at 4 °C in the dark. After washing, the cells were resuspended in FACS buffer with 0.1 μg/ml DAPI and filtered through a 50 μm cell strainer. Flow cytometry was done using BD LSR Fortessa, while FACS was performed with Sony SH100 Cell Sorter. Data were analysed using FlowJo (Tree Star).

### Isolation of gonadal hPGC by FACS

Human genital ridges were dissected in PBS and separated from the mesonephros followed by dissociation with TrypLE Express (Life Technologies) at 37°C for 20-40 minutes (depending on the tissue size) with pipetting every 5 minutes. Cells were washed, resuspended in FACS buffer (3% FBS and 5 mM EDTA in PBS) and incubated with anti-AP and anti-cKIT antibodies specified in Suppl. Table 5 for 15 minutes at room temperature with 10 rpm rotation in the dark. Cells were then washed in FACS medium and filtered through a 35 μm cell strainer. FACS was performed with Sony SH100 Cell Sorter and data were analysed using FlowJo (Tree Star). Cell populations of interest were sorted onto Poly-L-Lysine Slides (Thermo Scientific) and fixed in 4% PFA for IF analysis.

**Supplementary Table 1.** 4i medium composition.

| Component | Final Concentration | Supplier |
| --- | --- | --- |
| Knockout DMEM | - | GIBCO |
| Knockout serum replacement (KSR) | 20% | GIBCO |
| L-glutamine | 2 mM | GIBCO |
| Nonessential amino acids | 0.1 mM | GIBCO |
| 2-mercaptoethanol | 0.1 mM | GIBCO |
| Penicillin-Streptomycin | 100 U/ml (Penicillin) | GIBCO |
|  | 0.1 mg/ml (Streptomycin) |  |
| Human LIF | 20 ng/ml | Stem Cell Institute (SCI) |
| bFGF | 8 ng/ml | SCI |
| TGF-β1 | 1 ng/ml | Peprtech |
| CHIR99021 (CH) | 3 μM | Miltenyi Biotec |
| PD0325901 (PD) | 1 μM | Miltenyi Biotec |
| SB203580 (SB) | 5 μM | TOCRIS bioscience |
| SP600125 (SP) | 5 μM | TOCRIS bioscience |

**Supplementary Table 2.** hPGCLC medium composition.

| Component | Final Concentration | Supplier |
| --- | --- | --- |
| Advanced RPMI 1640 | - | GIBCO |
| B27 supplement | 1% | GIBCO |
| L-glutamine | 2 mM | GIBCO |
| Nonessential amino acids | 0.1 mM | GIBCO |
| Penicillin-Streptomycin | 100 U/ml (Penicillin) | GIBCO |
|  | 0.1 mg/ml (Streptomycin) |  |
| BMP2 | 500 ng/ml | SCI |
| Human LIF | 1 μg/ml | SCI |
| SCF | 100 ng/ml | R&D Systems |
| EGF | 50 ng/ml | R&D Systems |
| ROCK inhibitor (Y-27632) | 10 μM | TOCRIS bioscience |
| Poly-vinyl alcohol (PVA, 10%) | 0.25% | Sigma |

**Supplementary Table 3.**
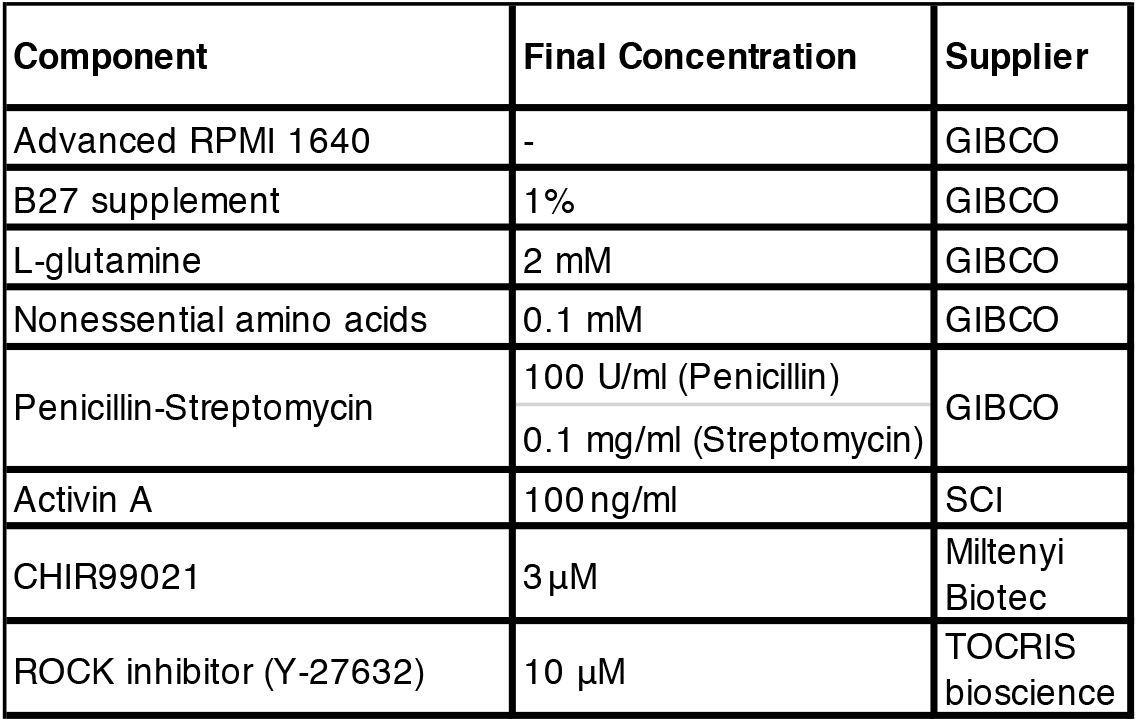
Mesendoderm (ME) medium composition.

**Supplementary Table 4.** Definitive endoderm (DE) medium composition.

| Component | Final Concentration | Supplier |
| --- | --- | --- |
| Advanced RPMI 1640 | - | GIBCO |
| B27 supplement | 1% | GIBCO |
| L-glutamine | 2 mM | GIBCO |
| Nonessential amino acids | 0.1 mM | GIBCO |
| Penicillin-Streptomycin | 100 U/ml (Penicillin)<br>0.1 mg/ml (Streptomycin) | GIBCO |
| Activin A | 100 ng/ml | SCI |
| BMP inhibitor (LDN193189) | 0.5 μM | Sigma |
| ROCK inhibitor (Y-27632) | 10 μM | TOCRIS bioscience |

### Immunofluorescence

For IF, hESCs were grown on ibiTreat 8-Well μ-Slides (Ibidi) on MEFs (GlobalStem), washed with PBS and fixed with 4% paraformaldehyde (PFA, Thermo Fisher Scientific) in PBS at room temperature (RT) for 10 minutes. This was followed by 3 washes in PBS and a 10-minute permeabilisation in 0.25% Triton X-100 (Sigma) in PBS. Next, the samples were incubated with the blocking buffer (0.1% Triton X-100, 5% normal donkey serum (NDS, Stratech) and 1% bovine serum albumin (BSA, Sigma)) for 30 minutes at RT. The samples were then incubated with primary antibodies (Suppl. Table 5) diluted in the blocking buffer overnight at 4 °C. The following day they were washed 3 times with the wash buffer (0.1% Triton X-100 in PBS) and incubated with Alexa fluorophore (AF-488, 568 and/or 647)-conjugated secondary antibodies (Invitrogen, used in 1:500 dilutions in the blocking buffer) specific for the host species of the primary antibodies for 1 hour at RT in the dark. After three washes in the wash buffer, the samples were incubated in PBS with 1 μg/ml DAPI for 10 minutes at RT. The samples were stored at 4 °C in the dark (up to 2 weeks) before imaging on Leica SP5 inverted confocal microscope. The images were analysed using Fiji software.

hPGCLC-containing EBs were fixed on D4 (unless otherwise specified) using 4% PFA in PBS at 4 °C for 1-2 hours. After two washes in PBS, the EBs were transferred to 10% sucrose solution in PBS and stored at 4 °C for 1 day. This was followed by a 1-day incubation in 20% sucrose in PBS at 4 °C. Finally, the EBs were embedded in OCT compound (CellPath), snap-frozen on dry ice and stored at −80 °C until cryosectioning. Cryorections of 8- m thickness were made on Superfrost Plus Micro slides (VWR) using a Leica Microsystems cryostat. Slides were stored at −80 °C or processed immediately. For IF, slides were air-dried at RT for 1 hour, washed in PBS 3 times 5 minutes each and permeabilised in 0.1% Triton X-100 in PBS for 30 minutes at RT. The cryosections on each slide were then circumscribed using ImmEdge Hydrophobic Barrier Pen (Vector Labs) and blocking solution (as above) was added for 30 minutes at RT. Samples were then incubated with primary (Suppl. Table 5) and secondary antibodies as specified for hESC IF above, but 1 g/ml DAPI was added to the secondary antibody mixtures. Finally, the slides were washed in the wash buffer twice and in PBS once and mounted with Prolong Gold Antifade Reagent (Molecular Probes). The images were acquired using Leica SP5 confocal microscope and analysed using Fiji software.

For fluorescence intensity quantification a Fiji plugin written by Dr. Richard Butler (Gurdon Institute, University of Cambridge, UK) was used. It counts 30-300 μm^2^ nuclei and measures fluorescence intensity in each channel. The data was then manually processed in Microsoft Excel to filter pluripotent cells (NANOG or OCT4 fluorescence intensity >50) where applicable and to calculate mean fluorescence intensities. Statistical analyses were performed using GraphPad Prism 7.

**Supplementary Table 5.**
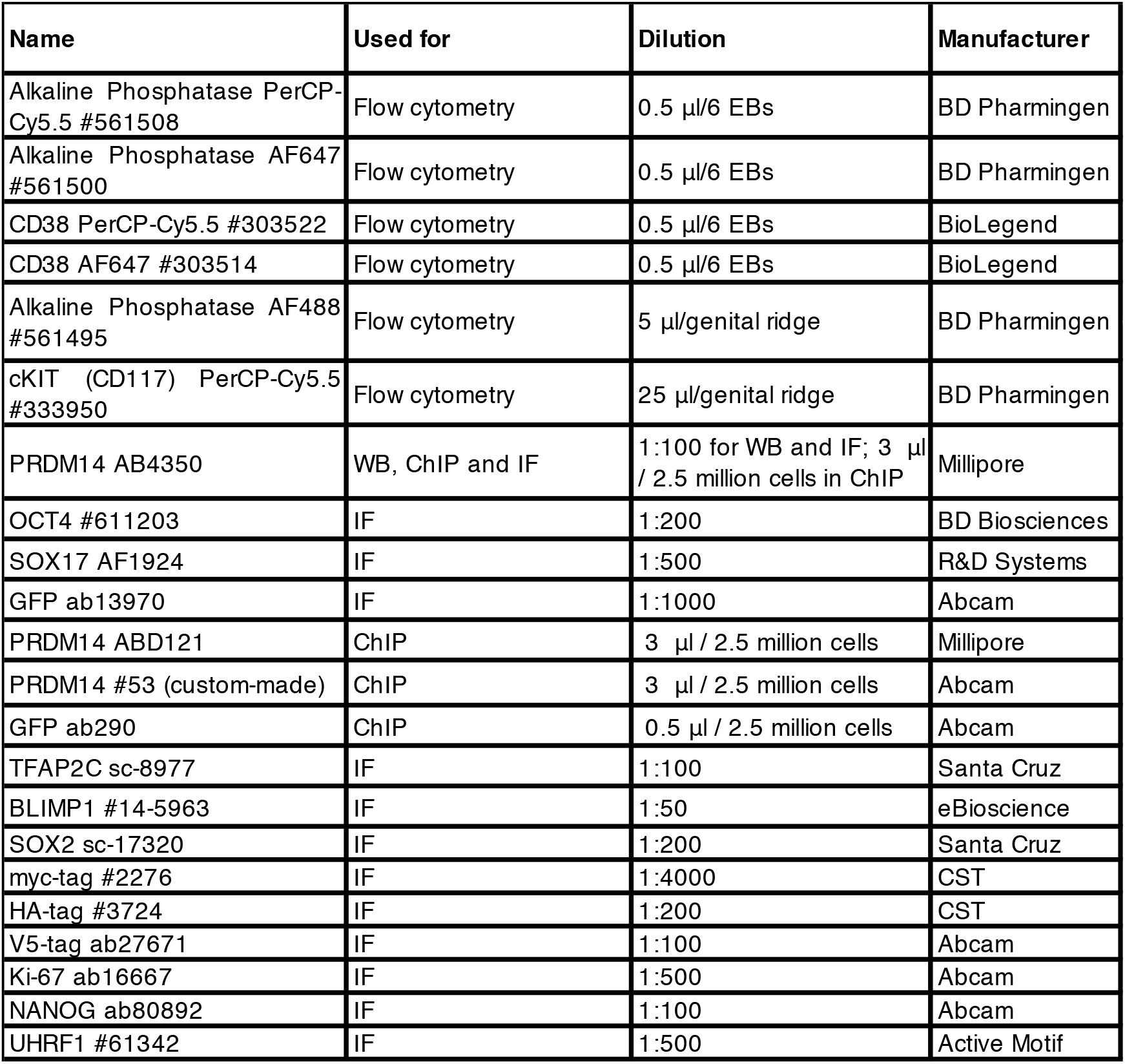
Antibodies used in the study.

### Plasmid constructions and cell line establishment

For gene targeting, CRISPR-Cas9 approach was used^5^. Annealed oligos (Suppl. Table 6) encoding corresponding gRNAs were ligated into pX330 vector5 digested with BbsI. gRNA sequences were chosen using MIT CRISPR tool (http://www.genome-engineering.org/crispr/) to be located close to the stop codon (to enable C-terminal fusions in PRDM14-T2A-Venus, PRDM14-AID-Venus, PRDM14-JAZ-Venus and SOX17-AID-Venus).

All other plasmids, including donor vectors for homology-directed repair were generated using In-Fusion cloning (Clontech) according to manufacturer’s recommendations, but scaling down the reaction to a total volume of 5 μl. Primers used for cloning are specified in Suppl. Table 6.

For knockins either electroporation or lipofection was used. For electroporation, ~250,000 trypsinised hESCs were resuspended in 600 μl PBS+Ca2++Mg2+ containing 50 μg gRNA plasmid and 50 μg homologous repair donor plasmid and electroporated in a 0.4-cm cuvette using Gene Pulser Xcell System (Bio-Rad) with a single 20 ms square-wave pulse (250 V). Lipofections were performed using 2 μg gRNA plasmid and 2 μg donor construct in OptiMEM medium (GIBCO) and Lipofectamin Stem reagent (Invitrogen) according to manufacturer’s recommendations. The volume of lipofectamine in μl was equal to the total amount of DNA in μg. For lipofection of PiggyBac transgenes a total of 1-5 μg DNA/100,000 cells was used, with the amount of PiggyBac transposase (PBase)-encoding plasmid equal to that of PiggyBac-delivered transgenes combined (in μg). Transfected cells were seeded onto drug-resistant DR4 MEFs (SCI) in 4i medium (ROCKi added for the first 24 hours; selection was initiated 48 hours after transfection).

After selection, individual clones were picked, expanded and genotyped using PCR. For this, genomic DNA was first extracted from cell pellets in the lysis buffer (10mM Tris pH 8.0, 100mM NaCl, 10mM EDTA and 0.5% SDS; Proteinase K (PK) was added immediately prior to lysis at a final concentration of 0.2 mg/ml) at 56 °C for 4 hours – overnight, followed by PK inactivation at 98 °C for 10 minutes. The supernatant was used directly in genotyping PCR performed with LongAmp (NEB) or PrimeStar GXL (Clontech) polymerase according to the manufacturers’ protocols. Primers used for genotyping are specified in Suppl. Table 6. In the case of transgenes expression, homogeneity and leakiness of induction was checked by IF.

For knockins (PRDM14-T2A-Venus, PRDM14-AID-Venus, PRDM14-JAZ-Venus and SOX17-AID-Venus) correct targeting in homozygous (by genotyping) clones was confirmed by Sanger sequencing. This was followed by the removal of the Rox-flanked puromycin resistance cassette by transient transfection with 1 μg of Dre-recombinase-encoding plasmid^6^. 2 days of hygromycin B selection (50 μg/ml) were followed by 2 days of negative selection using FIAU (200 nM)^7^. The obtained clones were again genotyped by PCR and sequenced to confirm antibiotic cassette excision. Venus intensity and homogeneity were then verified using Flow cytometry and IF.

For generation of cell lines that deplete PRDM14 upon IAA supplementation, AID-Venus was added at the C-terminus of PRDM14 using CRISPR (see above). This was followed by the addition of TIR1 transgene (pPB-CAG-OsTir1-myc-IRES-HygR or pPB-CAG-OsTir1-V5-T2A-Puro) by lipofection, along with pPBase. TIR1 cDNA and the 44-amino-acid AID sequence were kindly provided by Dr. Elphège Nora.

For generation of cell lines that deplete PRDM14 upon Cor supplementation, JAZ-Venus was added at the C-terminus of PRDM14 using CRISPR (see above). This was followed by the addition of COI1B transgene (pPB-CAG-HA-FboxOsTir1-OsCoi1b-IRES-HygR) by lipofection, along with pPBase. COI1B cDNA and the 43-amino-acid JAZ sequence were kindly provided by Dr. Ran Brosh.

For PRDM14-DD rescue, PRDM14-AID-Venus+TIR1 hESCs were co-lipofected with 0.05 μg pPB-CAG-myc-PRDM14-DD-IRES-HygR (low DNA amount was used to limit transgene copy number), 0.5 μg pPB-CAG-OsTir1-V5-T2A-Puro (to avoid TIR1 excision by PBase) and 0.55 μg PBase.

### Reverse-transcription quantitative PCR (qPCR)

Total RNA was extracted using RNeasy Mini Kit (QIAGEN) from unsorted cells or using Arcturus PicoPure (Thermo Fisher Scientific) from at least 1,000 sorted cells. cDNA was synthesized using QuantiTect Reverse Transcription Kit (QIAGEN). qPCR was performed on a QuantStudio 12K Flex Real-Time PCR machine (Applied Biosystems) using SYBR Green JumpStart Taq ReadyMix (Sigma) and specific primers (Suppl. Table 6). The ΔΔCt method was used for quantification of gene expression. Statistical analyses were performed using Microsoft Excel and/or GraphPad Prism 7.

### RNA-sequencing

RNA-seq was performed on PRDM14-AID-Venus competent 4i hESCs and hPGCLCs induced therefrom. Two biological replicates were used for each condition (Suppl. Table 7). For IAA-sensitive cells, two clones (cl11 and cl21) from the same hESC passage or the same hPGCLC induction were used as replicates. For the parental (“no TIR1”) control, the same cell line was used at different passages or inductions to yield two independent replicates. 10,000 AP+ 4i hESCs or 10,000 NANOS3-tdTomato+AP+ hPGCLCs (with the exception of hPGCLC cl21 replicate, where 3,000 cells were used) were sorted directly into 100 μl of extraction buffer from PicoPure RNA Isolation Kit (Applied Biosystems) for subsequent total RNA extraction according to manufacturer’s protocol. RNA was stored at −80 °C and its quality and quantity were checked by Agilent RNA 6000 Pico Kit with Bioanalyzer (Agilent Technologies) and Qubit (Thermo Fisher Scientific). RNA-seq library was prepared from 10 ng input RNA using the end-to-end Trio RNA-seq library prep kit (Nugen) following the manufacturer’s protocol but omitting the AnyDeplete step. In short, the protocol contains the following steps: DNAse treatment to remove DNA from RNA; first strand and second strand cDNA synthesis to produce the reverse complement of the input RNAs; cDNA purification using Agencourt AMPure XP beads (Backman Coulter); single primer isothermal amplification (SPIA) to stoichiometrically amplify cDNAs; enzymatic fragmentation and end repair; sequencing adaptor (index) ligation; product purification using AMPure beads; library amplification (4 cycles were used); library purification using AMPure beads. Libraries were then quantified by qPCR using NEBNExt Library Quant Kit (NEB) for Illumina on QuantStudio 6 Flex Real-Time PCR System (Applied Biosystems). Fragment size distribution and the absence of adapter dimers was checked using Agilent TapeStation 2200 and High Sensitivity D1000 ScreenTape. Finally, RNA-seq libraries were subjected to single-end 50 bp sequencing on HiSeq 4000 sequencing system (Illumina). 24 indexed libraries (including samples from this work and 8 others) were multiplexed together and sequenced in two lanes of a flowcell, resulting in >30 million reads per sample.

**Supplementary Table 7.** Summary of RNA-seq samples. Note that two different IAA-sensitive clones (cl11 and cl21) were used as replicates. The samples of the parental cell line (Par) lacking TIR1 were obtained in two replicates (reps).

| 4i hESCs | hPGCLCs |
| --- | --- |
| Par no IAA (2 reps) | Par no IAA (2 reps) |
| Par +100μM IAA (2 reps) | Par +100μM IAA (2 reps) |
| Cl11 no IAA | Cl11 no IAA |
| Cl11 +100μM IAA | Cl11 +100μM IAA |
| Cl21 no IAA | Cl21 no IAA |
| Cl21 +100μM IAA | Cl21 +100μM IAA |
| 8 samples | 8 samples |

### RNA-seq analysis

The library sequence quality in demultiplexed fastq files was checked by FastQC (v0.11.5)^9^ and the low-quality reads and adaptor sequences were removed by Trim Galore (v0.4.1)^10^ using the default parameters. The pre-processed RNA-seq reads were mapped to the human reference genome (UCSC GRCh38/hg38) using STAR (2.6.0a)^11^ (--outFilterMismatchNoverLmax 0.05 --outMultimapperOrder Random -- winAnchorMultimapNmax 100 --outFilterMultimapNmax 100) guided by the ENSEMBL (Release 87) gene models. Read counts per gene were extracted using TEtranscripts^12^ and normalized by DEseq2 in R^13^. Differential expression analysis was also performed using DEseq2. The resulting gene expression table (Suppl. Table 9) was used for downstream analyses in Microsoft Excel and R. Pearson’s correlation analysis was performed using the R cor command. Unsupervised hierarchical clustering was performed using the R hclust function with (1-Pearson’s correlation coefficient) as distance measures. The R prcomp function was used for Principal component analysis (PCA). Gene set enrichment analysis was performed using the GSEA software by the Broad Institute^14^. Gene Ontology (GO) analysis was performed using DAVID^15^.

For comparative differential expression analysis of PRDM14 depletion in hPGCLCs and TFAP2C and PRDM1 KOs from^16^ the samples were identically pre-processed and mapped to the human reference genome (UCSC GRCh38/hg38), sequencing reads were re-normalised together using DEseq2 to generate a new integrated gene expression table (Suppl. Table 11). A similar procedure was performed for comparisons with mouse Prdm14 KO from^17^, where processed reads were mapped to the mouse reference genome (UCSC GRCm38/mm10), and high-confidence human-mouse “one-to-one” orthology assignments for protein-coding genes were obtained from ENSEMBL (Release 87) (Suppl. Table 13). Venn diagrams were plotted using VennPainter^18^ and p-values for two overlapping datasets were calculated using a generalized hypergeometric test for multiple samples^19^. For the co-expression analysis of differentially expressed genes, the gene-based Pearson’s correlation coefficients were calculated for PRDM14-AID-Venus competent hESC and hPGCLC samples using R. All pairwise correlations r < 0.8 between genes were removed. The matrix was imported as an adjacency matrix into the R igraph package^20^, and the Louvain method for community detection was performed on the resulting graph. For comparison with microarray data^21^, the published data was normalized, log2-transformed and the differential expression was evaluated using the R limma package^22^.

### Chromatin immunoprecipitation (ChIP) and ChIP-seq

ChIP was performed using SimpleChIP Enzymatic Chromatin IP Kit (with Magnetic Beads) from Cell Signalling Technology (CST) following manufacturer’s recommendations with modifications. Recipes for all buffers are provided in Suppl. Table 8. The following samples were included in the analysis (one replicate per condition): 4i hESCs (cl11 no IAA), 4i+IAA hESCs (cl11+IAA) and hPGCLCs (cl11 and cl21 pooled; no IAA). For each ChIP 2.5 million unsorted 4i hESCs or 4 million unsorted hPGCLCs (D4 EBs containing 60-70% PGCLCs as assessed by flow cytometry) were used. The cells were washed in PBS, filtered through 50 μm strainer and fixed in 1% formaldehyde at RT for 10 min. Formaldehyde was quenched by adding 450 μl of 10x glycine at RT for 10 min, followed by centrifugation at 500 g, 4 °C for 5 minutes. The cells were then washed twice in ice-cold PBS with protease inhibitor cocktail (PIC) and the pellets were snap frozen on dry ice and stored at −80 °C. To prepare nuclei, cells were thawed on ice, washed in ice-cold buffers A and B, resuspended in 100 μl buffer B and transferred to a thin-walled Diagenode 1.5 ml tubes. Next, 0.35 μl micrococcal nuclease (MNase) was added and the suspension was incubated for 23 minutes at 37 °C, shaking at 800 rpm to enzymatically digest the chromatin. The addition of 10 μl 0.5M EDTA for 2 minutes on ice was used to stop the digestion. Next, the nuclei were pelleted at 13,000 rpm, 4 °C for 1 minute, resuspended in 100 μl of ChIP buffer 1 and kept on ice for 10 minutes, followed by sonication (3x 30 seconds on/off, high output) using Bioruptor sonicator (Diagenode). The lysate was cleared by centrifugation at 10,000 rpm, 4 °C for 10 minutes and transferred to a fresh DNA LoBind 1.5 ml tube (Eppendorf). For chromatin immunoprecipitation, 400 μl ChIP buffer 2 was added for a total of 500 uL/sample; 10 μl were removed and stored at −80 °C to serve as 2% input. Antibodies were then added to each immunoprecipitation (IP) reaction (0.5 μl anti-GFP (ab290, abcam); 3 μl anti-PRDM14 (ABD121, Millipore), 3 μl anti-PRDM14 (ABD4350, Millipore); 3 μl anti-PRDM14 (#53 custom-made by abcam)) and IPs were incubated overnight rotating at 10 rpm, 4 °C.

Equal volumes of Protein A and Protein G-conjugated magnetic beads (Invitrogen) were combined (for a total of 20 μl per IP) in a DNA LoBind 1.5 ml tube, washed in 200 μl blocking buffer, and incubated with 500 μl blocking buffer for 1 hour rotating at 10 rpm, 4 °C. The tubes were placed on a magnetic rack (Diagenode) to remove the supernatant and another 1-hour incubation was performed. Then the supernatant was removed, the beads were resuspended in 20 μl blocking buffer and stored at 4 °C for up to 24 hours until the use in ChIP.

After overnight IP, 20 μl of pre-blocked Protein A/G beads were added to each sample, followed by rotation at 10rpm, 4 °C for 2 hours. The tubes were placed on a magnetic rack and the supernatant was removed. The beads were then washed three times with 1 ml low-salt buffer (each wash involved a 5-minute rotation at 10rpm, 4 °C), once with high-salt buffer and once with LiCl buffer. Finally, the beads were resuspended in 50 μl elution buffer and transferred to 0.2 ml PCR stripes. The inputs were thawed on ice and supplemented with 50 μl elution buffer. All samples were incubated at 65 °C, 1100 rpm for 30 minutes to elute the protein-DNA complexes from the beads. The stripe was then placed onto a magnetic rack and the eluate was transferred into a fresh 0.2 ml PCR stripe to which 2 μl 5M NaCl and 2 μl proteinase K (10 mg/ml) were added. The samples were then incubated at 65 °C, 1100 rpm for 2 hours for decrosslinking.

Finally, for the use in ChIP-qPCR, DNA was purified using MinElute columns (QIAGEN) following the manufacturer’s protocol. Alternatively, for ChIP-seq library preparation, DNA was purified using Agencourt AMPure XP beads (Beckman Coulter) as follows. 1.8x volume of AMPure beads were mixed with each sample, incubated for 10 minutes to bind DNA to the beads and washed 2×30 seconds with 200 μl 80% ethanol using a magnetic rack. The beads were air-dried for 5-10 minutes to reach complete ethanol evaporation. 33 μl of elution buffer (10 mM Tris-HCl, pH 8.0 – 8.5) was added and incubated with the beads for 2 minutes to elute the ChIP DNA. The PCR stripe was placed on the magnetic rack and 30 μl of eluate was transferred into fresh tubes.

For ChIP-seq libraries were prepared using KAPA HyperPrep Kit following the manufacturer’s instructions. Briefly, the protocol contains the following steps: end-repair and A-tailing; sequencing adapter (index) ligation; product purification using AMPure beads; library amplification using KAPA real-time Library Amplification Kit (11 cycles were used for ChIP libraries and 7 for input libraries to achieve similar concentration range); product purification using AMPure beads. Libraries were then quantified by qPCR using NEBNExt Library Quant Kit for Illumina (NEB) on QuantStudio 6 Flex Real-Time PCR System (Applied Biosystems). Fragment size distribution and the absence of adapter dimers was checked using Agilent TapeStation 2200 and High Sensitivity D1000 ScreenTape. Finally, ChIP-seq libraries were subjected to single-end 50 bp sequencing on HiSeq 4000 sequencing system (Illumina). 8 indexed libraries were multiplexed in one lane of a flowcell, resulting in >20 million reads per sample.

**Supplementary Table 8.** Buffer recipes for ChIP.

| **Buffer A** | **1x** |
| --- | --- |
| Buffer A stock [μl] | 250 |
| H2O [μl] | 750 |
| DTT [μl] | 0.5 |
| PIC [μl] | 5 |

| **Buffer B** | **1x** |
| --- | --- |
| Buffer B stock [μl] | 275 |
| H2O [μl] | 825 |
| DTT [μl] | 0.6 |

| **ChIP buffer 1** | **1x** |
| --- | --- |
| ChIP buffer stock [μl] | 10 |
| H2O [μl] | 90 |
| PIC [μl] | 0.5 |

| **ChIP buffer 2** | **1x** |
| --- | --- |
| ChIP buffer stock [μl] | 40 |
| H2O [μl] | 360 |
| PIC [μl] | 2 |
| tRNA (Sigma) [10mg/ml] [μl] | 2 |

| **Elution** | **1x** |
| --- | --- |
| Elute buffer | 25 |
| H2O | 25 |

| **Blocking buffer** | **1x** |
| --- | --- |
| ChIP buffer (CST) [μl] | 150 |
| H2O [μl] | 1110 |
| PIC (Roche) [μl] | 75 |
| BSA (Sigma) (10%) [μl] | 150 |
| tRNA (Sigma) [10mg/ml] [μl] | 15 |

| **Low salt** | **1x** |
| --- | --- |
| ChIP buffer | 300 |
| H2O | 2700 |

| **High salt** | **1x** |
| --- | --- |
| ChIP buffer | 100 |
| H2O | 900 |
| NaCl | 70 |

| **LiCl Wash buffer** | **For 250 mL** |
| --- | --- |
| 20mM Tris-HCl pH 8.0 | 5 ml of 1M |
| 2mM EDTA | 1 ml of 0.5M |
| 250 mM LiCl | 2.65 g |
| 1% NP-40 | 2.5 ml |
| 1% Deoxycholate | 2.5 g |
| H2O | 241.5 ml |

### ChIP-seq analysis

The library sequence quality in demultiplexed fastq files was checked by FastQC (v0.11.5)^9^ and the low-quality reads and adaptor sequences were removed by Trim Galore (v0.4.1)^10^ using the default parameters. The trimmed ChIP-seq reads were aligned to the human reference genome (UCSC hg38) by the BurrowsWheeler Aligner (0.7.15-r1142-dirty)^23^. Samtools (version: 1.3.1)^24^ was used to remove unmapped and low mapping quality reads (options: view -F 4 -q 20). Subsequently, duplicated reads and reads mapped to unlocalized contig (random), unplaced contigs (ChrUn) and blacklisted regions (obtained from http://mitra.stanford.edu/kundaje/akundaje/release/blacklists/hg38-human/) were removed using Samtools rmdup. This was followed by preliminary peak calling using macs2 callpeak^25^ against the corresponding inputs (options: -g 3e9 --keep-dup all). Percentage of reads in peaks was calculated using featureCounts^26^ and visualised with multiQC^27^. Before peak calling, to compare the number of peaks between the three high-quality samples, the library size for all libraries was downsampled to 25 million reads. Peaks were then called again using macs2 callpeak against the corresponding inputs (options: -g 3e9 --keep-dup all --nomodel – extsize 157). The resultant peak files of the three samples were merged to generate a consensus set of 6486 peaks using bedtools merge (options: -d 100)^28^. The consensus peaks were processed and clustered by DeepTools computeMatrix and plotHeatmap^29^. Peak annotation, motif enrichment and differential peak analyses were performed by HOMER^30^. DeepTools bamCoverage (options: -extendReads 157 --binSize 10 -- normalizeUsing CPM --ignoreForNormalization chrX chrY) was used to generate bigwig file for visualization in Integrative Genomics Viewer^31^. The integrated ChIP-seq and RNA-seq table (Suppl. Table 12) was generated using R to correlate changes in gene expression to PRDM14 binding. Venn diagrams were plotted using the R/Bioconductor ChIPseeker package^32^.

---

**Supplementary Figure 1.**
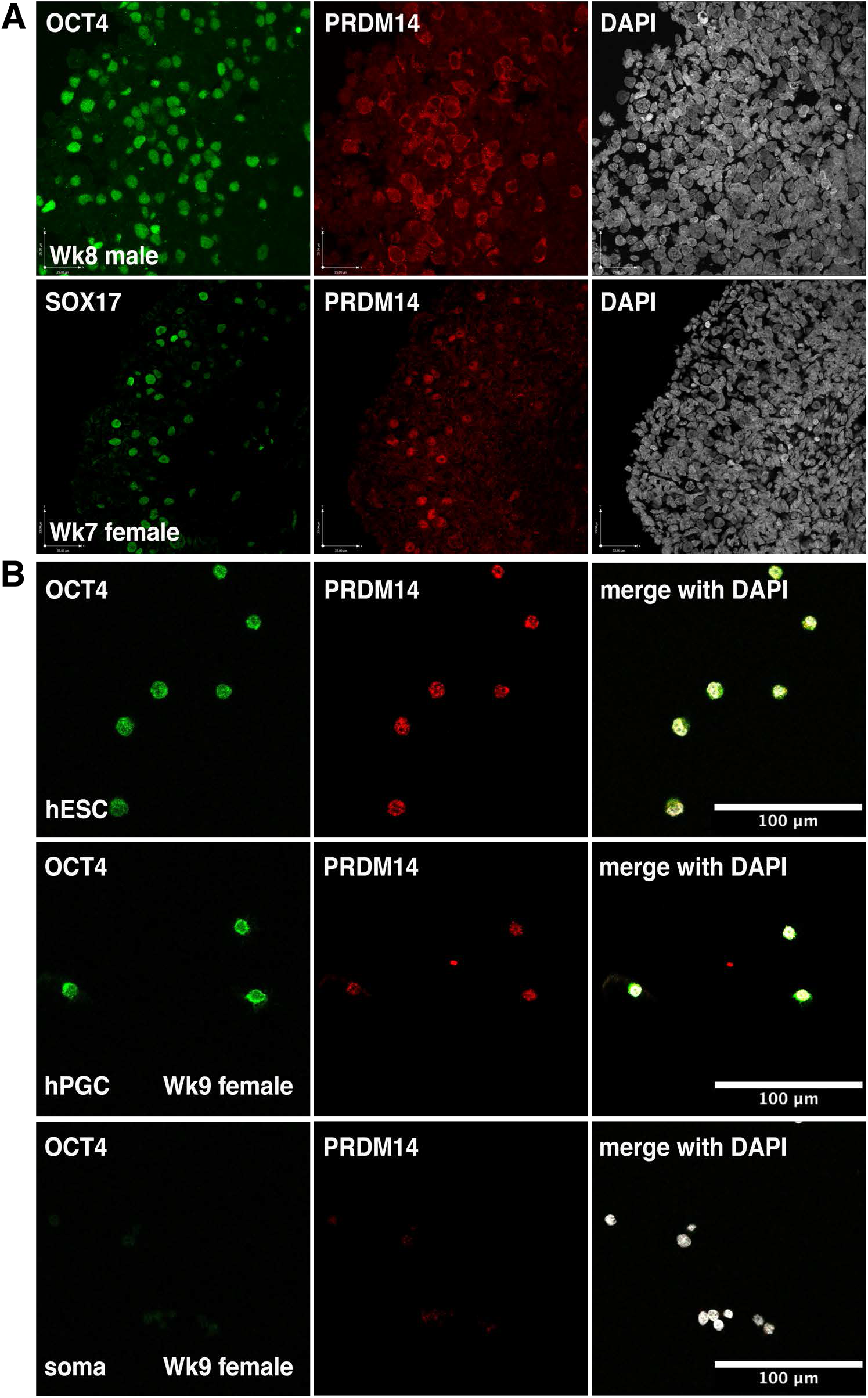
Nuclear PRDM14 expression in human gonadal PGCs. (**A**) Representative IF images of PRDM14 staining using two different antibody batches in Wk7 and Wk8 human female gonadal sections. hPGCs were marked by OCT4 or SOX17 expression. Nuclei were counterstained by DAPI. Note both nuclear and cytoplasmic PRDM14 in the upper sample (Wk8 male). (**B**) IF on sorted AP+ hESCs and AP+cKIT+ gonadal hPGCs. hPGCs and pluripotent hESCs were marked by OCT4. Nuclei were counterstained by DAPI. AP^-^cKIT^-^ population (gonadal soma) was used as a negative control. Similar results were obtained for a Wk9 male embryo. Scale bar is 100 μm.

**Supplementary Figure 2.**
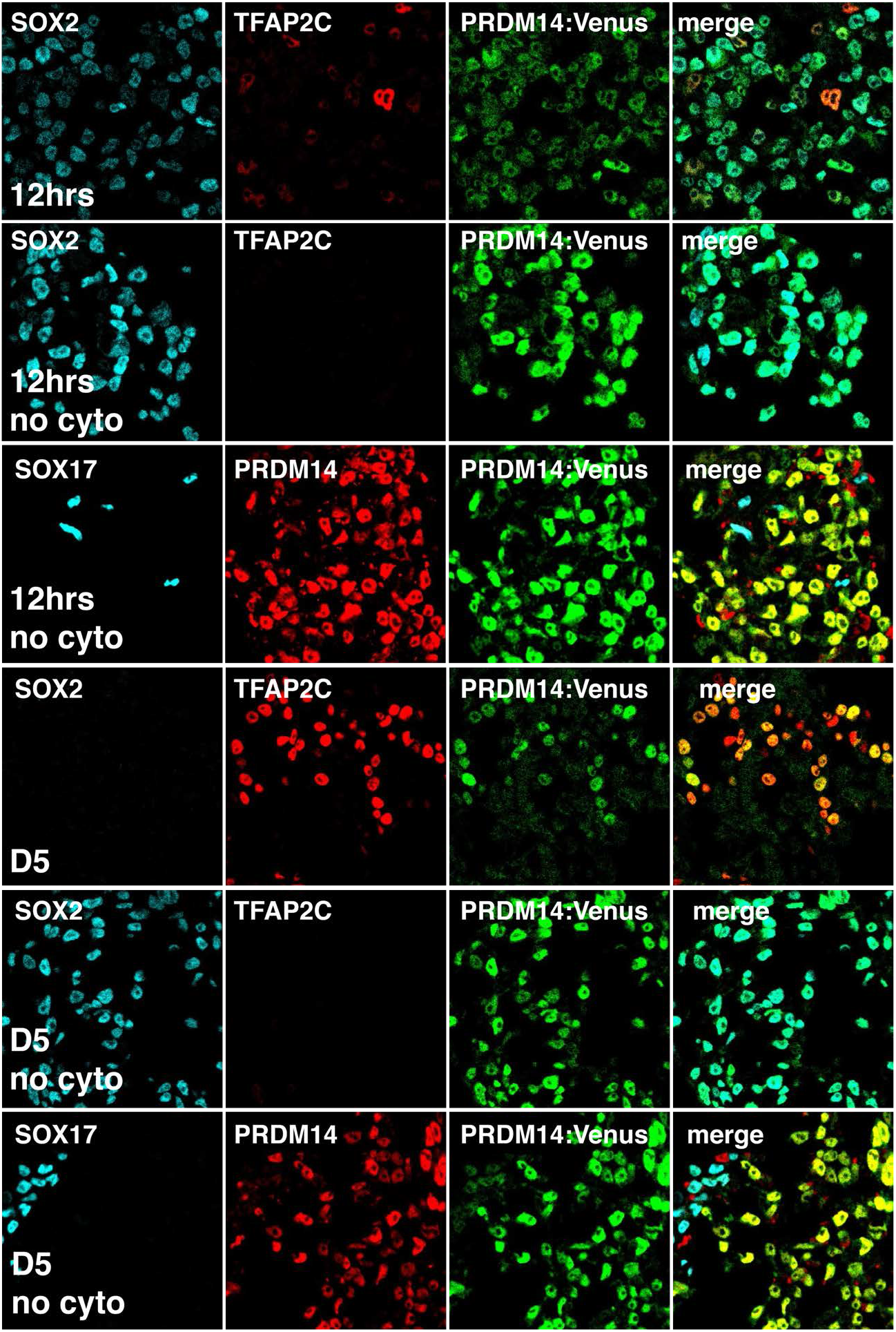
PRDM14-Venus and SOX2 expression during PGCLC specification. IF analysis of 12hrs and D5 EBs differentiated from PRDM14-AID-Venus fusion reporter cell line with or without cytokines (“+cyto” and “no cyto”, respectively). Expression of PRDM14 and SOX2 is shown; hPGCLCs in “+cyto” EBs are marked by SOX17 or TFAP2C. Note that “no cyto” differentiation in the basal medium yields no hPGCLCs and SOX17+ cells likely belong to other lineages.

**Supplementary Figure 3.**
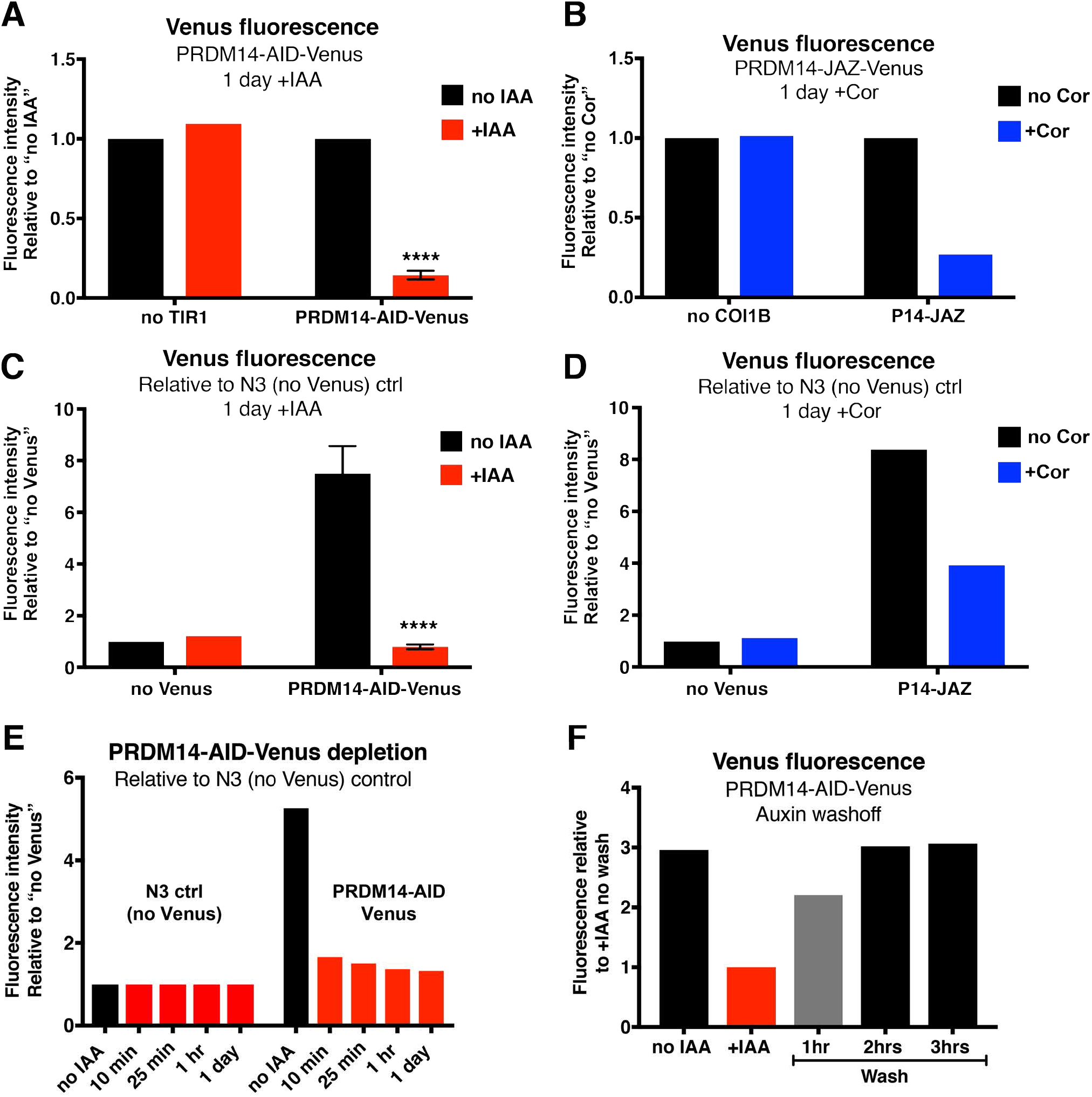
Venus fluorescence quantification shows higher efficiency of AID compared to JAZ degron. (**A**) Related to Fig.3B. PRDM14-AID-Venus cells were grown with or without IAA for 1 day. To focus on pluripotent cells that would normally express PRDM14, fluorescence intensities were measured in the nuclei of NANOG-positive cells. Data show the mean of n=4 (2 independent experiments using 3 clones), **** P<0.0001 (twoway ANOVA followed by Sidak’s multiple comparison test) and n=1 for “no TIR1” control. A cell line lacking TIR1 hormone receptor (“no TIR1”) was used as a negative control for PRDM14-AID-Venus depletion. (**B**) Related to Fig.3C. PRDM14-JAZ-Venus cells were grown with or without Cor for 1 days. To focus on pluripotent cells that would normally express PRDM14, fluorescence intensities were measured in the nuclei of OCT4-positive cells. A cell line lacking COI1B hormone receptor (“no COI1B”) was used as a negative control for PRDM14-JAZ-Venus depletion. (**C**) Data from (A) normalised to a control cell line lacking AID-Venus KI. (**D**) Data from (B) normalised to a control cell line lacking JAZ-Venus KI. (**E**) Kinetics of Venus fluorescence depletion in PRDM14-AID-Venus hESCs. hESCs were treated with IAA for indicated time periods. Homogeneous depletion with Venus intensity close to “no Venus control” was reached within 25 minutes, as indicated by IF staining in Fig.3B. (**F**) Kinetics of Venus fluorescence restoration in PRDM14-AID-Venus hESCs. hESCs were treated with IAA for 2 days, washed and grown without IAA for indicated periods of time.

**Supplementary Figure 4.**
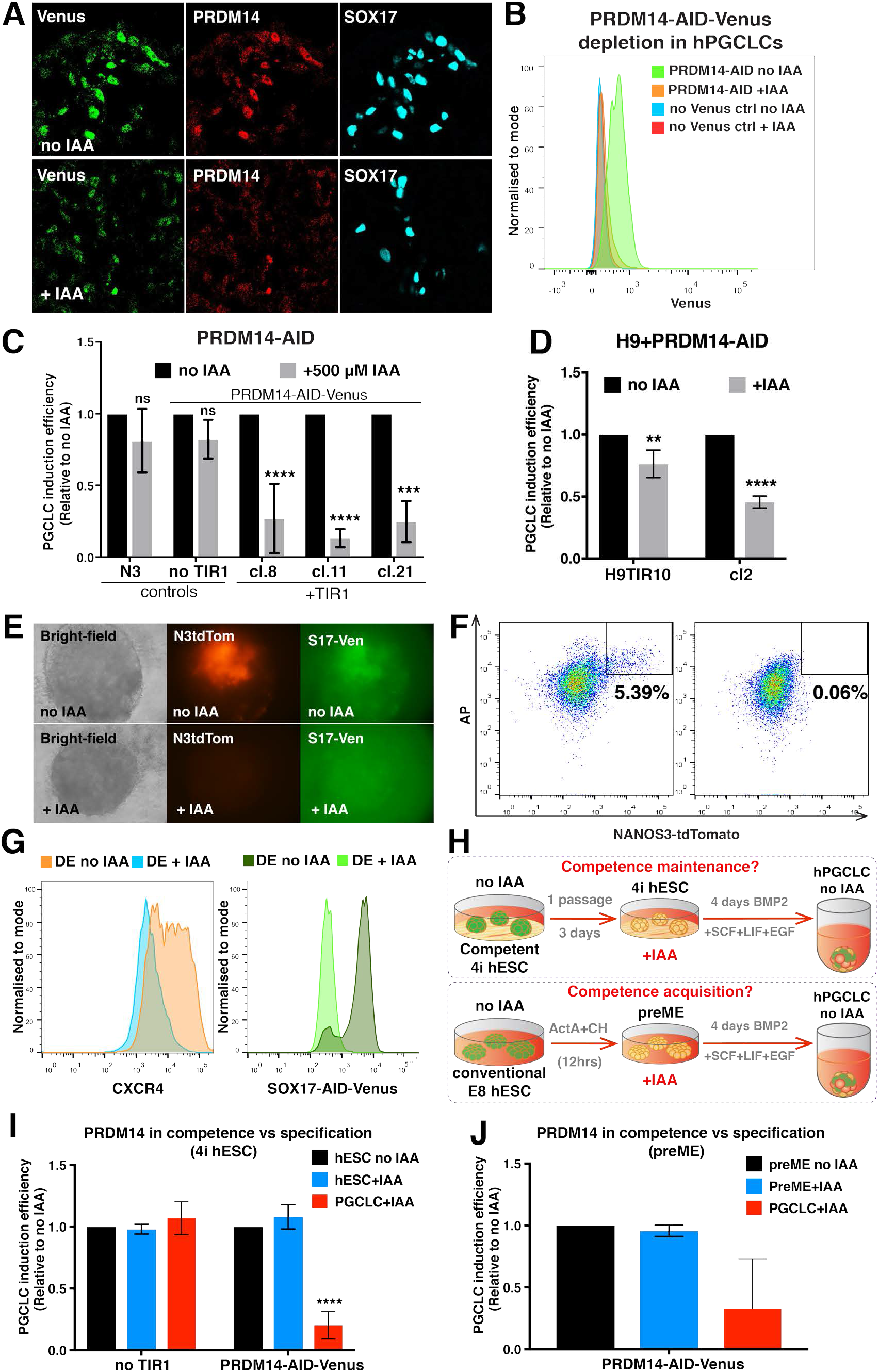
AID system allows studying the roles of PRDM14 and SOX17 TFs in cell fate decisions. (**A**) If analysis of PRDM14-AID-Venus depletion in D4 IAA-treated EBs. SOX17 marks hPGCLCs. (**B**) Representative flow plots showing Venus fluorescence in AP+CD38+ hPGCLCs induced with or without auxin. (**C**) hPGCLC induction efficiencies from PRDM14-AID-Venus differentiated with or without 500 μM IAA concentrations (note a different concentration compared to Fig.4B). The induction efficiency of untreated cells (“no IAA”) were set to 1. Data show mean+/-SD for n=3-5 per clone; ns – not significant, **** P<0.0001 (Two-way AnOVA followed by Sidak’s multiple comparison test). (**D**) PRDM14 depletion using AID in H9 hESC background also reduces hPGCLC specification. Note that this cell line lacks NANOS3-tdTomato reporter and hPGCLCs were identified by AP and CD38 staining. H9TIR10 is a parental cell line expressing TIR1 but lacking the AID-Venus KI. Data show mean+/-SD for n=3 independent experiments per clone, ** P<0.01, **** P<0.0001 (Two-way ANOVA followed by Sidak’s multiple comparison test). (**E**) Epifluorescence photographs show colocalization of NANOS3-tdTomato and SOX17-AID-Venus fluorescence in D4 EBs; both signals are reduced upon IAA treatment. (**F**) Representative flow cytometry plots showing hPGCLC induction using the SOX17-AID-Venus cell line with or without IAA. (**G**) SOX17 degradation decreases definitive endoderm (DE) specification. Representative flow cytometry plots; DE cells are marked by CXCR4 expression. (**H**) Experimental workflow of PRDM14 depletion in competent hESCs, as well as during preME induction. PRDM14 role in competence maintenance was assessed by depleting it in competent (4i) hESCs, followed by washing off the IAA for PRDM14 restoration and inducing hPGCLCs (without IAA). To test if PRDM14 is required for competence acquisition, it was depleted during the generation of the preME competent cells. Auxin was subsequently removed and hPGCLCs were induced without IAA. (**I**) PRDM14-AID depletion in competent hESCs does not reduce hPGCLC induction therefrom. IAA was added for 1 passage (3 days) prior to hPGCLC induction and washed of before the onset of hPGCLC differentiation. hESCs grown without auxin were differentiated with or without IAA as above. Data show mean+/-SD for n=2-4 independent experiments per clone, **** P<0.0001 (Two-way ANOVA followed by Sidak’s multiple comparison test). (**J**) PRDM14-AID depletion during the acquisition of hPGC-competent state via pre-mesendoderm (preME) does not reduce hPGCLC induction therefrom. IAA was added for 12 hours during preME generation from conventional hESCs and washed off before the onset of hPGCLC differentiation. preME induced without auxin was differentiated with or without IAA as above. Data show mean+/-SD for n=2 independent experiments (2 clones).

**Supplementary Figure 5.**
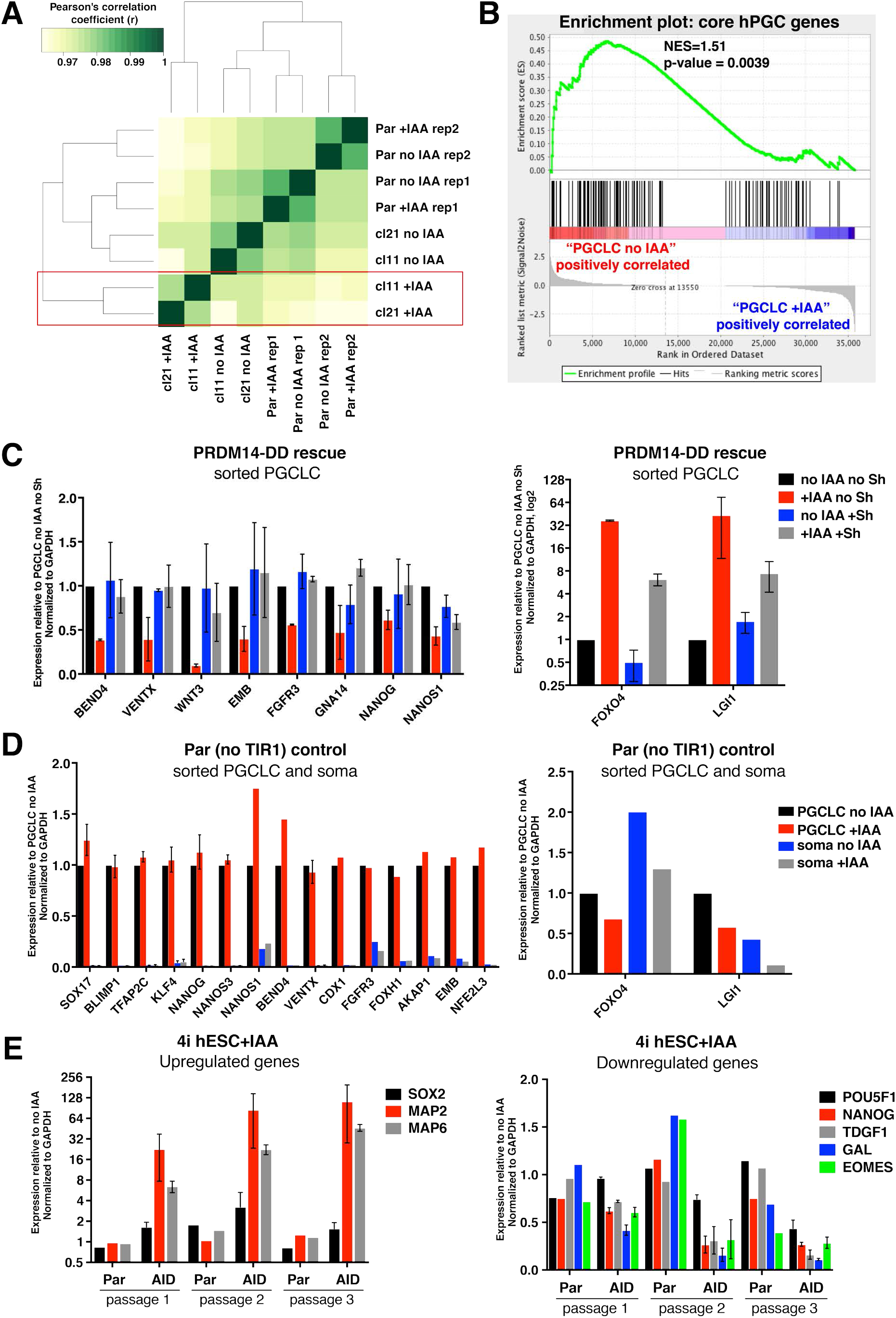
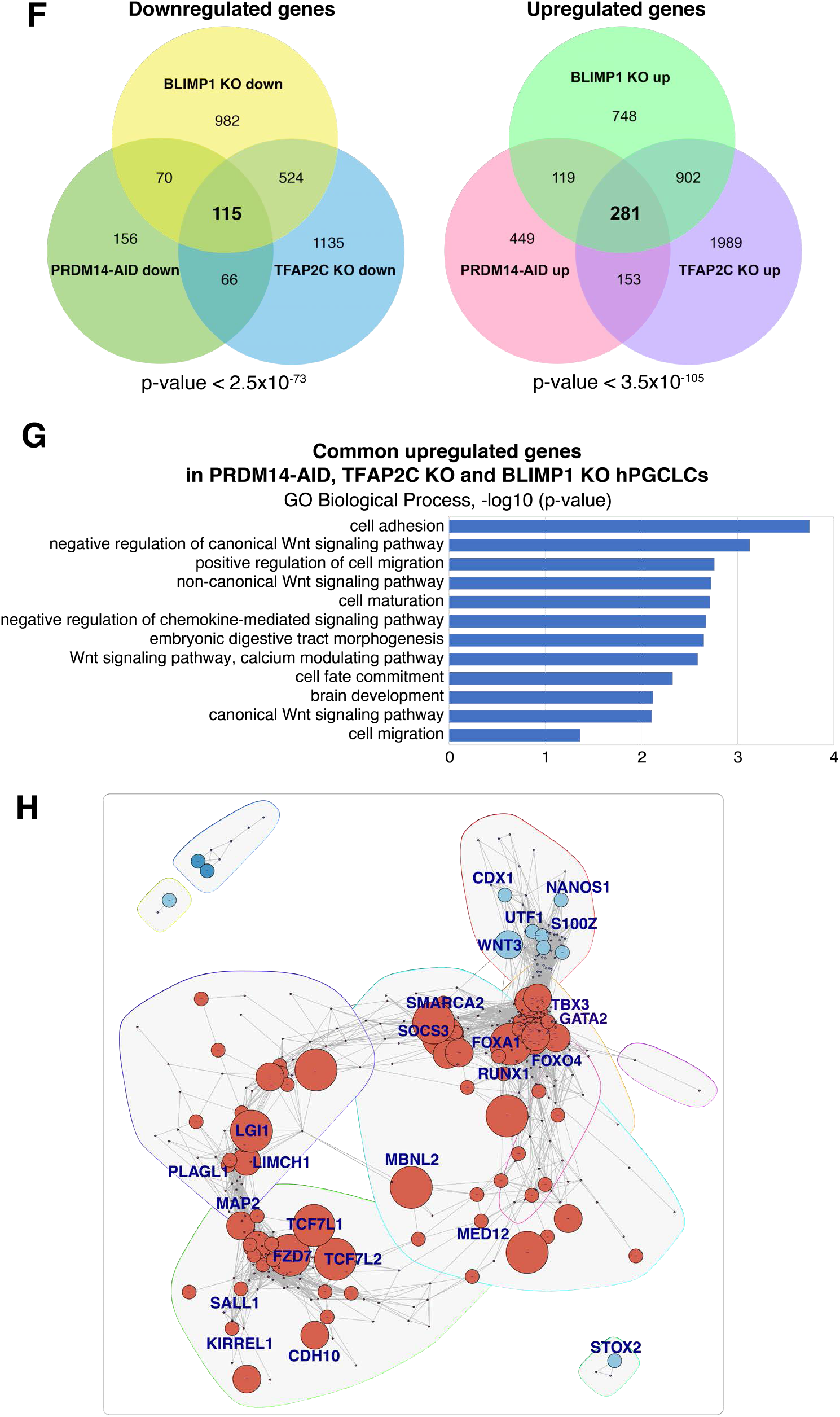
Transcriptional changes in hPGCLCs induced without PRDM14. (**A**) Heatmap showing Pearson’s correlation between hPGCLC RNA-seq samples. “Par” denotes parental control lacking TIR1 and hence unresponsive to auxin; “r” – Pearson’s correlation coefficient. PRDM14-depleting clones cluster separately and are highlighted in red. (**B**) Gene set enrichment analysis (GSEA) of 123 hPGC-specific genes on the transcriptome of hPGCLCs specified with or without IAA. PRDM14-deficient hPGCLCs (“PGClC+IAA”) are negatively correlated with the gene set, if compared against “PGCLC no IAA” control. (**C**) qPCR shows transcriptional rescue of tested DEGs by ectopic PRDM14-DD. hPGCLCs were differentiated with or without IAA and/or Shield-1 (Sh) and sorted as NANOS3-tdTomato+AP+ cells; soma denotes the double-negative population from the same experiments. Data show gene expression relative to no IAA no Sh PGCLCs and normalized to *GAPDH,* mean+/-SEM of n=2 independent experiments. (**D**) Parental (“no TIR1”) control for qPCR validation of RNA-seq (see Fig.6G). hPGCLCs were sorted as NANOS3-tdTomato+AP+ cells, while soma denotes the double-negative population from the same experiments. Data show gene expression relative to no IAA hPGCLCs and normalized to *GAPDH,* mean+/-SEM of n=2-5 independent experiments, * P<0.05, ** P<0.01, *** P<0.001, **** P<0.0001 (multiple t-tests). (**E**) qPCR on unsorted PGC-competent (4i) PRDM14-AID hESCs cultured with or without IAA for 3 consecutive passages. The parental cell line lacking TIR1 is shown as a control. Data show mean+/-SEM of n=2 hormone-sensitive clones; n=1 for the parental control. Note downregulation of pluripotency genes and upregulation of pro-neuronal differentiation transcripts. Transcriptional changes in hPGCLC induced without PRDM14. (**F**) Venn diagrams highlighting significant overlap between transcriptional changes in hPGCLCs lacking PRDM14, TFAP2C or BLIMP1 (relative to corresponding WT controls). P-values were obtained using a modified hypergeometric test (Polanski et al., 2014). (**G**) GO analysis performed on shared upregulated genes. The graph shows top non-redundant GO terms as -log10(p-value). (**H**) Co-expression network of differentially expressed genes in hPGCLCs (+IAA vs no IAA, log2FC > 1.5, p-value < 0.05). Nodes were scaled according to the number of PRDM14 ChlP-seq peaks assigned to the gene. The network was partitioned into overlapping communities using the Louvain method (see Methods). Representative direct targets of PRDM14 were labelled. Upregulated genes were highlighted in orange and downreguated in blue. Note that downregulated PRDM14 targets form a separate co-expression cluster.

**Supplementary Figure 6.**
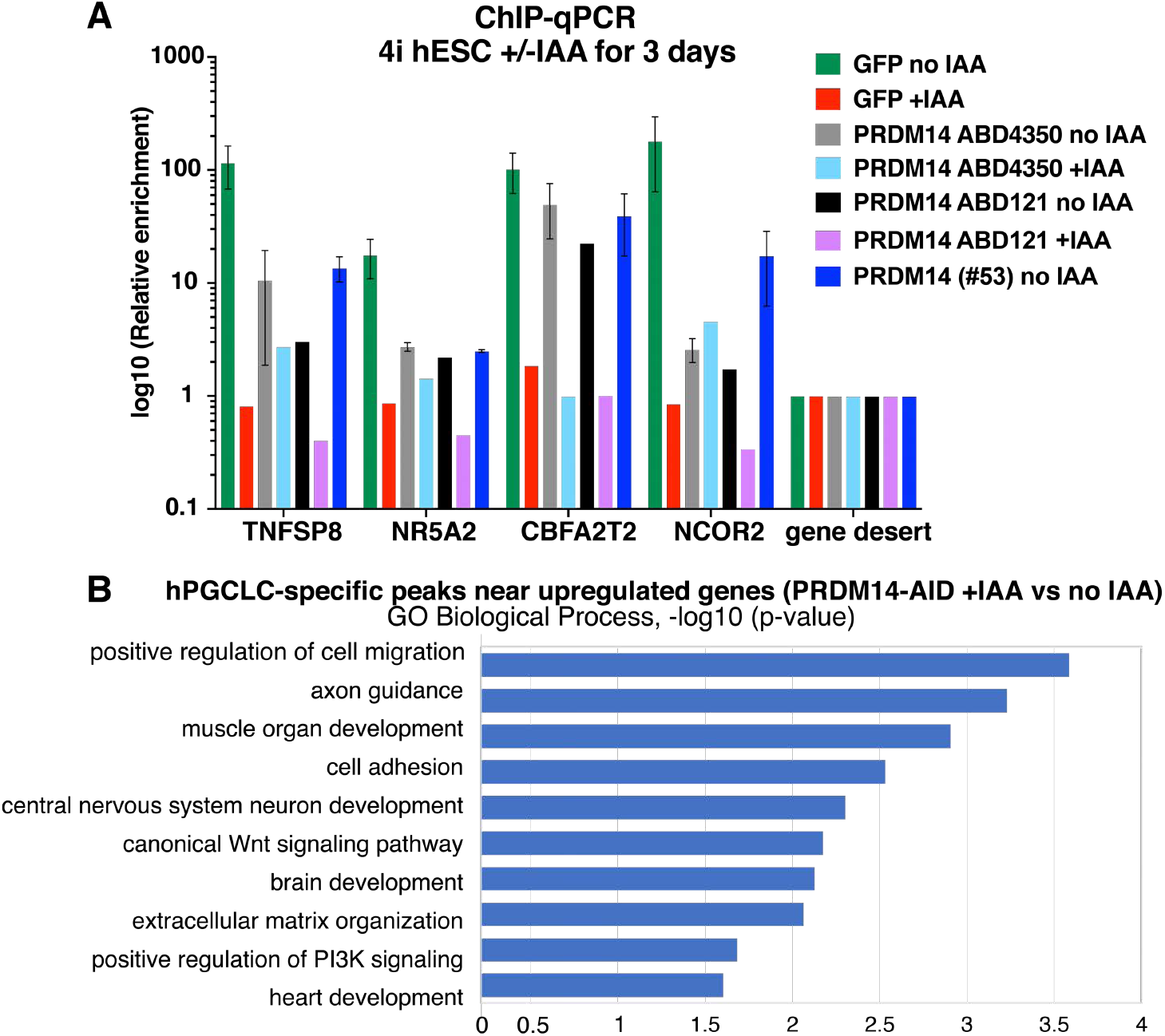
PRDM14-Venus ChIP optimisation and GO analysis on direct targets of PRDM14. (**A**) ChlP-qPCR in competent PRDM14-AID-Venus hESCs using different antibodies. Anti-GFP antibody allows higher enrichment than the tested anti-PRDM14 antibodies. The primers were designed to span selected PRDM14 ChlP-seq peaks in conventional hESCs from (Chia et al. 2010). Data show log10 enrichment normalised to input and relative to a gene desert region (control), mean+/-SEM for n=1-3 independent experiments. (**B**) GO analysis on hPGCLC-specific PRDM14 peaks (q-value<0.05) near genes upregulated in hPGCLCs upon PRDM14 depletion. Top non-redundant GO terms are shown as –log10(p-value).

**Figure 7.**
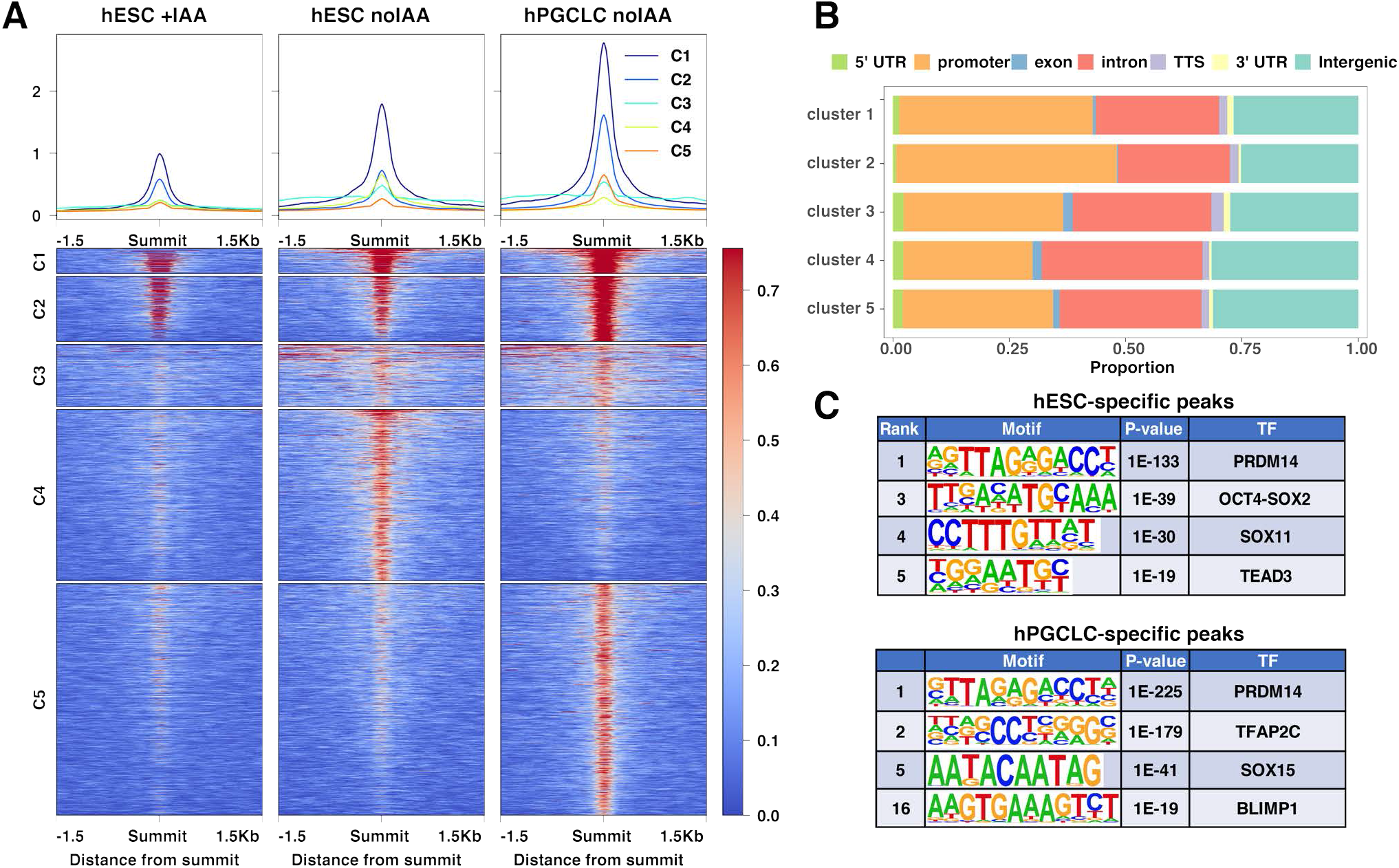
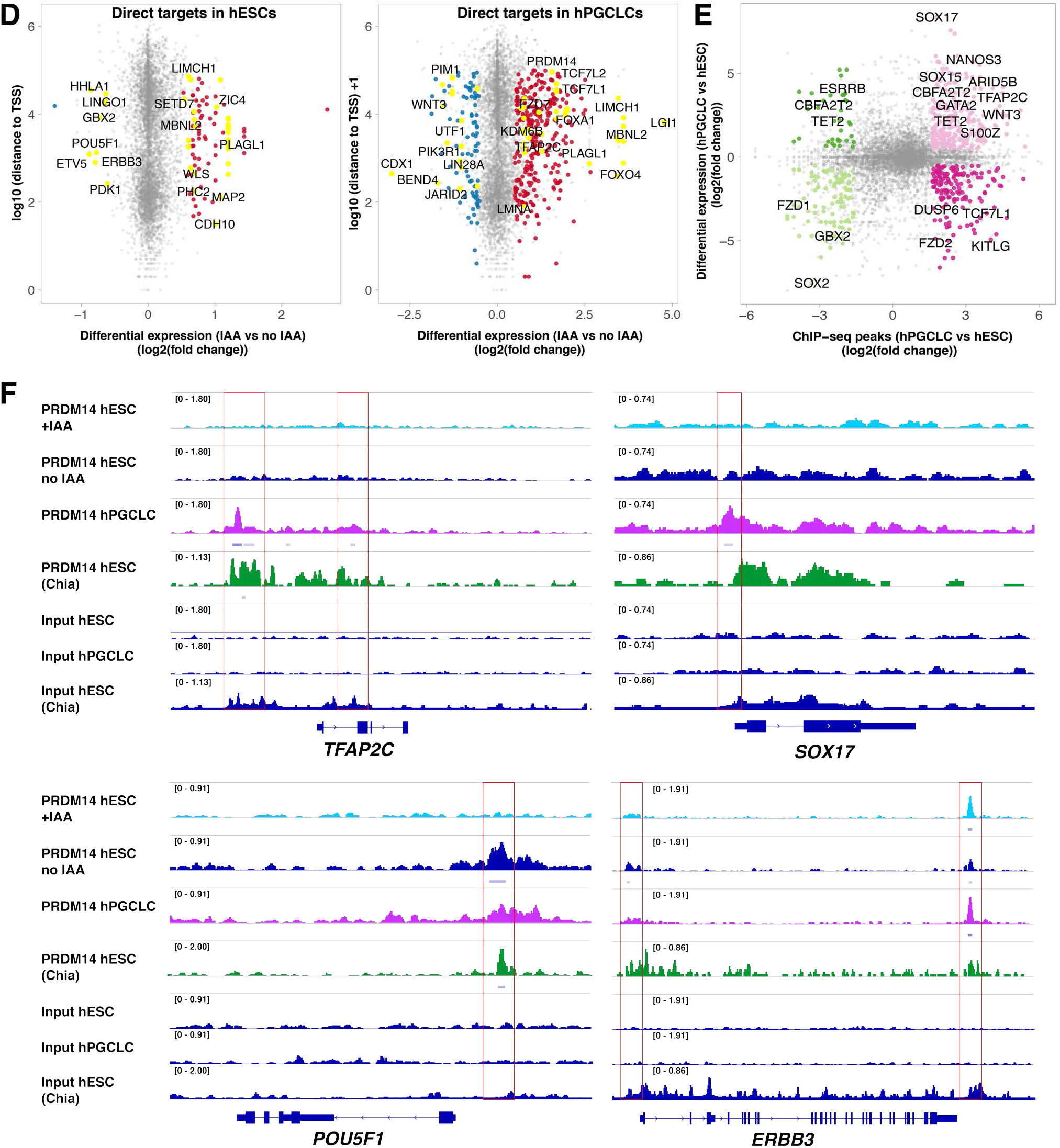
ChIP-seq analysis shows hESC- and hPGCLC-specific PRDM14 binding patterns. (**A**) Consensus PRDM14-bound peaks (6486) are separated into 5 clusters by *k*-means clustering. The ChIP signals (counts per million per 10-bp bin) of each peak are shown in the clustered heatmaps (bottom panel). The profile plots (top panel) show the average ChIP signals of each cluster. (**B**) Distribution of clustered PRDM14 peaks in the genome. Note that PRDM14 predominantly binds to promoters. (**C**) Top ranking motifs identified by Homer *de novo* motif analysis within hESC- and hPGCLC specific PRDM14 peaks (FC>3, p-value<0.0001). ChIP-seq analysis shows hESC- and hPGCLC- specific PRDM14 binding patterns. (**D**) Scatter plots highlighting examples of direct PRDM14 targets in hESCs and hPGCLCs. Direct targets were defined as genes with at least one PRDM14 ChIP-seq peak (q-value <0.05) within 100kb of the TSS and showing significant changes in expression upon IAA treatment (log2FC<(−0.5) or >0.5 and padj<0.05). (**E**) PRDM14 binds to regulatory regions of key hPGC-related genes. The scatter plot shows differential PRDM14 peaks (FC>3, p-value<0.0001) correlated to genes differentially expressed between hESCs and hPGCLCs (log2FC<(−0.5) or >0.5 and padj<0.05). (**F**) Examples of loci bound by PRDM14 in hPGCLCs and hESCs. Note that the peak downstream of *ERBB3* is preserved in hESCs after IAA treatment. PRDM14 peaks in indicated samples were visualized using IGV software. PRDM14 ChIP-seq dataset from (Chia et al., 2010) was added for comparison.

**Supplementary Figure 8.**
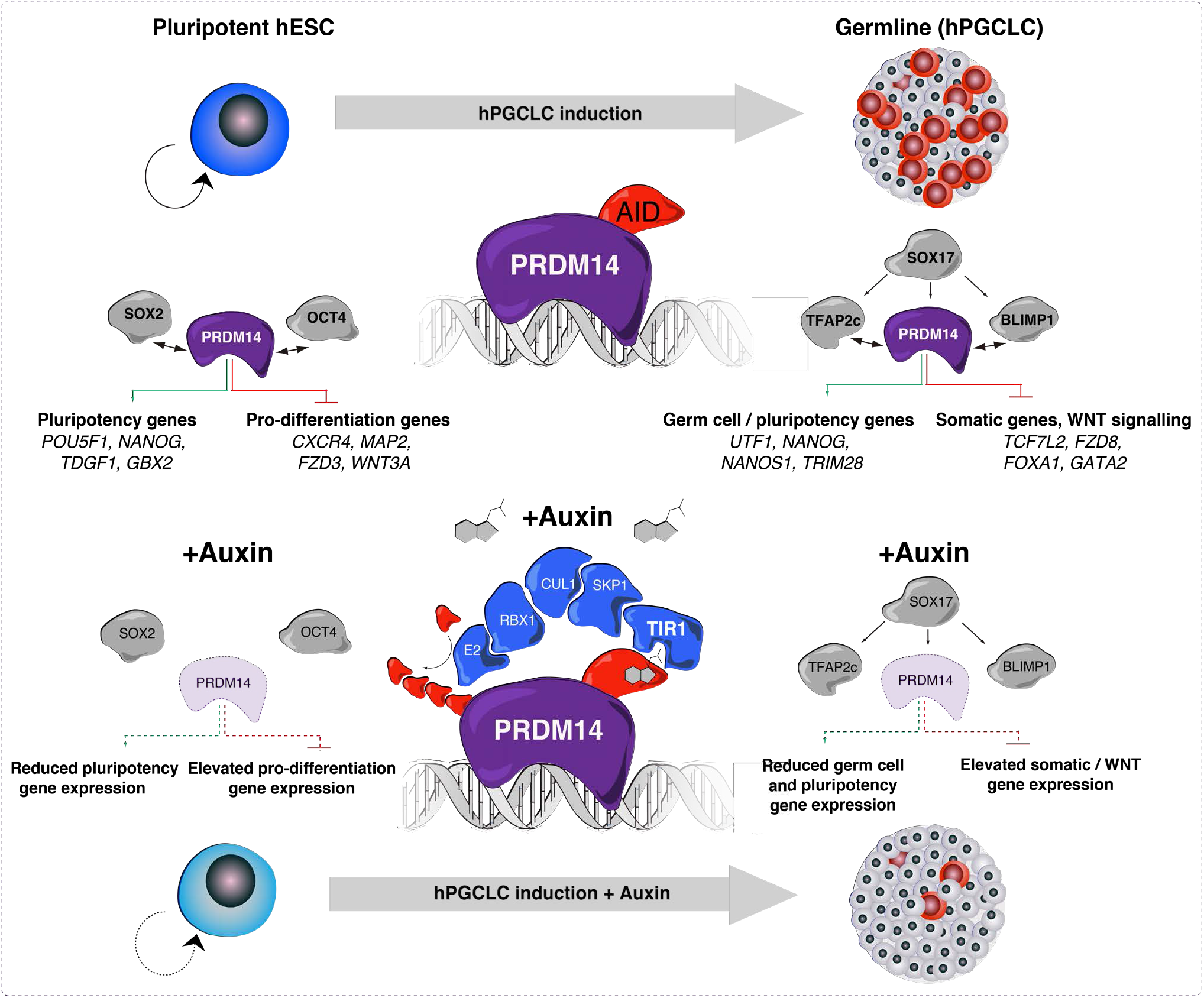
Graphical summary of key findings. Model summarising the context-dependent PRDM14 roles in human pluripotency and hPGCLC specification. Using PRDM14-AID-Venus we were able to map PRDM14 binding in hESCs and hPGCLCs, as well as deplete the protein with high time resolution. This showed that PRDM14 regulates target genes through promoter binding together with SOX2/OCT4 in hESCs or TFAP2C and BLIMP1 in hPGCLCs. PRDM14, TFAP2C and BLIMP1 are all downstream of SOX17. In hESCs, PRDM14 activates the expression of pluripotency genes and represses pro-differentiation genes. Similarly, in hPGCLCs, PRDM14 represses somatic transcripts, but activates germ cell- and pluripotency-related genes. Representative examples of activated and repressed genes are shown. When auxin is added PRDM14-AID fusion protein is recruited to the E3 ubiquitin ligase via an F-box auxin receptor TIR1. It leads to rapid protein ubiquitination and degradation, which allows time-resolved functional studies.

